# Molecular and spectroscopic characterization of green and red cyanine fluorophores from the Alexa Fluor and AF series

**DOI:** 10.1101/2020.11.13.381152

**Authors:** Christian Gebhardt, Martin Lehmann, Maria M. Reif, Martin Zacharias, Thorben Cordes

## Abstract

The use of fluorescence techniques has had an enormous impact on various research fields including imaging, biochemical assays, DNA-sequencing and medical technologies. This has been facilitated by the availability of numerous commercial dyes, but often information about the chemical structures of dyes (and their linkers) are a well-kept secret. This can lead to problems for applications where a knowledge of the dye structure is necessary to predict (unwanted) dye-target interactions, or to establish structural models of the dye-target complex. Using a combination of spectroscopy, mass spectrometry and molecular dynamics simulations, we here investigate the molecular structures and spectroscopic properties of dyes from the Alexa Fluor (Alexa Fluor 555 and 647) and AF series (AF555, AF647, AFD647). Based on available data and published structures of the AF and Cy dyes, we present two possible structures for Alexa Fluor 555. We also resolve conflicting reports on the linker composition of Alexa Fluor 647. A comprehensive comparison between Alexa Fluor and AF dyes by continuous-wave absorption and emission spectroscopy, quantum yield determination, fluorescence lifetime and anisotropy spectroscopy of free and protein-attached dyes, supports the suggestion that the Alexa Fluor and AF dyes have a high degree of structural similarity. In addition, we compared Alexa Fluor 555 and Alexa Fluor 647 to their structural homologs AF555 and AF(D)647 in single-molecule FRET applications. Both pairs showed excellent performance in solution-based smFRET experiments using alternating laser excitation demonstrating that the AF-fluorophores are an attractive alternative to Alexa- and Cy-dyes for smFRET studies, and suggesting their usefulness for other fluorescence applications.

## 1. Introduction

The exploitation of fluorescence techniques has had a large impact on various research fields and specific applications such as optical imaging, biochemical assays, DNA-sequencing, and medical technologies. The molecular contrast agents, i.e., the light absorbing and emitting molecules used, and their properties govern the success of these applications (for instance in PCR-based amplification of disease-related genomes^1^) and the information depth of state-of-the-art techniques in specialized research fields such as single-molecule^2^ and super-resolution microscopy^3–9^. Whereas fluorescent proteins dominate live-cell applications, in most other settings where high photostability and tailored functional properties are required^2,10^, synthetic organic fluorophores predominate.

The common molecular scaffolds of modern synthetic organic fluorophores are fluoresceins, rhodamines, carbon- and silicon-pyronines, rylenes, bodipys, and cyanines^10^. They all feature intense absorption and emission in the visible spectrum^10^. Years of structural optimization has resulted in commercially available compounds with favorable photophysical properties and reactive linkers for bioconjugation. The general structural design of such commercial fluorophores aims at high absorbance cross sections, high fluorescence quantum yields, and low rates for internal conversion and intersystem crossing e.g., achieved by the exclusion of heavy atoms to reduce the latter (see e.g., ref. ^2^ and references cited therein). In addition, self-healing^11–16^, self-blinking^17^, and photoactivatable dyes^18^, fluorescent sensors for ions^19^ and pH etc. have become (commercially) available. Small-molecule additives^20–25^ are frequently used as intermolecular reaction partners for dyes to either improve their performance by reduction of photodamage by triplet-states^9^, oxygen^9,26^ and other reactive fluorophore species, or to achieve photoswitching^8,27^.

The increasing availability of dyes from commercial sources has been a boon to research and medicine. Companies, however, have often not been forthcoming with information on the chemical structures of the dyes (and their linkers), which has been an obstacle for some applications. Prominent examples are the nucleic acids stains of the SYBR family (SYBR Green, SYBR Gold)^28^, the Alexa Fluor dye series (Alexa Fluor 555)^29^, and the ATTO dye family (ATTO643). A few structures from these suppliers have recently been made available (SYBR Green^30^, ATTO647N^31^, ATTO655^32^). There are many applications where knowledge of the chemical structure of a dye is not necessary, e.g., when using Alexa Fluor 555 in imaging^33–41^ and spectroscopic studies^42–45^. However, Alexa Fluor 555 and Alexa Fluor 647 are dyes that are frequently and successfully used for single-molecule FRET in combination with other fluorophores^46–50^, or as a donor-acceptor pair^51–56^. They have become a popular choice because of their favorable performance in many assays, and this is largely due to their high water solubility and the absence of strong (unwanted) interactions between dye and target following conjugation.

For Alexa Fluor 647, the chemical structure is known, but there are conflicting reports in the literature on the linker length connecting the two sulphonated SO_3_^-^ groups (both 3-carbon^47,57,58^ or 4-carbon atoms^59–63^ have been reported), as well as the structure of the maleimide-linker connecting the chromophore to a biological target. For Alexa Fluor 555, on the other hand, there is no verified information on its fluorophore class or molecular structure. Fluorescent lifetimes similar to Cy3^16^ suggested that it might have a cyanine core^57^, and a chemical structure was proposed but never verified experimentally^62,64–66^. The lack of unequivocal structural data for these fluorophores limits their proper use for FRET-restrained structural modelling, and in situations where an understanding of dye-target interactions are important^67^, such as in molecular dynamics simulations^56,62,68^.

We have here studied the molecular and spectroscopic properties of Alexa Fluor 555 and Alexa Fluor 647 in relation to other cyanine fluorophores with known molecular structures (Cy3, sulfo-Cy3, AF555, Cy5, sulfo-Cy5, AF(D)647, Figure 1).

**Figure 1.**
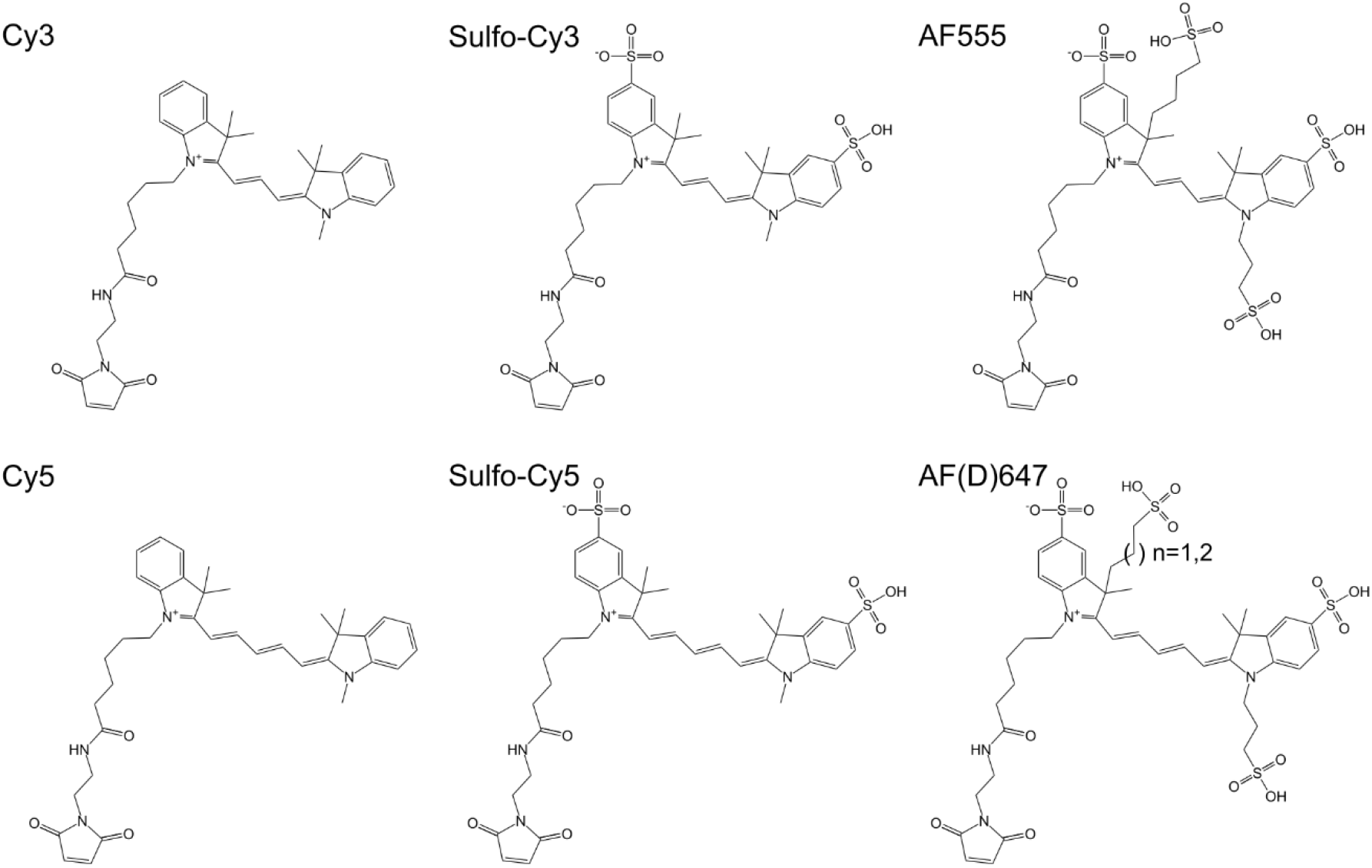
Chemical structures of published and characterized cyanine fluorophores from the Cy-series and the AF-fluorophores. The AF-fluorophore homologue of Cy5 is available in two versions AFD647 with n=1 and AF647 with n=2.

Using a combination of visible spectroscopy, mass spectrometry, and molecular dynamics simulations, we show that Alexa Fluor 555 has a cyanine-based fluorophore core identical to Cy3. Based on the available data, we present two possible structures for Alexa Fluor 555, and suggest the most likely one to be similar to that of AF555 (structure of AF555 see Figure 1). We further clarified the precise molecular structure of commercial Alexa Fluor 647 maleimide to settle contradicting reports on its linker composition. Using a combination of spectroscopic techniques including continuous-wave absorption and emission spectroscopy, quantum yield determination, fluorescence lifetime and anisotropy spectroscopy of free and protein-attached dyes, we compared Alexa Fluor 555 and Alexa Fluor 647 to their structural homologs Cy3/AF555 and Cy5/AF(D)647 in single-molecule FRET applications. Based on the high similarity of the molecular and spectroscopic parameters presented in this manuscript, we explored and characterized the performance of donor-acceptor pairs AF555-AFD647 for smFRET in direct comparison to Alexa Fluor 555-Alexa Fluor 647. Both dye pairs showed good performance in solution-based smFRET experiments using alternating laser excitation. This suggests that the AF-fluorophores are an attractive alternative to Alexa- and Cy-dyes for smFRET studies and that they should perform well in other fluorescence applications.

## 2. Material and Methods

### Sample preparation and labelling of proteins

MalE single and double cysteine variants were obtained and fluorophore-labelled as described previously^69,70^. The cysteine positions for fluorophore attachment were chosen based on the open and closed x-ray crystal structures of MalE (1OMP, 1ANF, respectively). The double cysteine variants were (i) stochastically labelled with the maleimide derivative of the dyes Alexa Fluor 555 and Alexa Fluor 647 (ThermoFischer Scientific, A20346 & A20347), and AF555, AFD647 & AF647 (Jena Bioscience, APC-007, APC-009 and APC-009) for FRET measurements. (ii) Corresponding single cysteine variants were labelled with one fluorophore as indicated. His-tagged proteins were incubated in buffer containing 1 mM DTT to keep all cysteine residues in a reduced state. Subsequently proteins were immobilized on a Ni Sepharose 6 Fast Flow resin (GE Healthcare). The resin was incubated 2-4 h at 4°C with 25 nmol of each fluorophore dissolved in labelling buffer 1 (50 mM Tris-HCl pH 7.4-8.0, 50 mM KCl, 5% glycerol) and subsequently washed sequentially with 1 CV labelling buffer 1 and 2 (50 mM Tris-HCl pH 7.4-8.0, 50 mM KCl, 50% glycerol) to remove unbound fluorophores. Bound proteins were eluted with 500 ml of elution buffer (50 mM Tris-HCl pH 8, 50 mM KCl, 5% glycerol, 500 mM imidazole) The labelled protein was further purified by size-exclusion chromatography (ÄKTA pure, Superdex 75 Increase 10/300 GL, GE Healthcare) to remove remaining fluorophores and aggregates. For all proteins, the labelling efficiency was higher than 80% (Supplementary Figure S1) and donor-acceptor pairing at least 30 % for double-cysteine variants (Supplementary Figure S10).

### Sample handling for quantum-yield, time-resolved anisotropy, and single-molecule FRET measurements

The labelled MalE proteins were stored in 50 mM Tris-HCl pH 7.4, 50 mM KCl with 1 mg/ml bovine serum albumin (BSA) at 4°C for less than 7 days. The samples were stored at protein concentrations between 100-500 nM and diluted for the measurements indicated as described below.

### Single-molecule FRET measurements and data analysis

ALEX experiments were carried out by diluting the labelled proteins to concentrations of ≈50 pM in 50 mM Tris-HCl pH 7.4, 50 mM KCl supplemented with the ligand maltose as described in the text and figures. Before each experiment, the coverslip was passivated for 5 minutes with a 1 mg/ml BSA solution in PBS buffer. The measurements were performed without photostabilizer, which showed little effects on the resulting data quality (Supplementary Figure S2), which is in contrast to the pair Cy3B/ATTO647N used previously for amino-acid binding proteins^52,71,72^ and ribosome recycling factor ABCE1^73^ where the addition of TX/MEA had a significant positive impact.

Data acquisition and correction procedures were performed for confocal measurements similar to the procedure as described by Hellenkamp *et al*.^47^. Solution based smFRET experiments were performed on a homebuilt confocal ALEX microscope as described in ^74^. All samples were studied using a 100 μl drop of buffer (50 mM Tris-HCl pH 7.4-8.0, 50 mM KCl) on a coverslip. The fluorescent donor molecules were excited by a diode laser at 532 nm (OBIS 532-100-LS, Coherent, USA) operated at 60 μW at the sample in alternation mode (50 μs alternating excitation and a 100 μs alternation period). The fluorescent acceptor molecules were excited by a diode laser at 640 nm (OBIS 640-100-LX, Coherent, USA) operated at 25 μW at the sample. The lasers were combined and coupled into a polarization maintaining single-mode fiber (P3-488PM-FC-2, Thorlabs, USA). The laser light was guided into an epi-illuminated confocal microscope (Olympus IX71, Hamburg, Germany) by a dual-edge beamsplitter ZT532/640rpc (Chroma/AHF, Germany) and focused to a diffraction-limited excitation spot by a water immersion objective (UPlanSApo 60x/1.2w, Olympus Hamburg, Germany). The emitted fluorescence was collected through the same objective, spatially filtered using a pinhole with 50 μm diameter and spectrally split into donor and acceptor channel by a single-edge dichroic mirror H643 LPXR (AHF). Fluorescence emission was filtered (donor: BrightLine HC 582/75 (Semrock/AHF), acceptor: Longpass 647 LP Edge Basic (Semroch/AHF)) and focused onto avalanche photodiodes (SPCM-AQRH-64, Excelitas). The detector outputs were recorded by a NI-Card (PCI-6602, National Instruments, USA).

Data analysis was performed using a home written software package as described in ^70^. Single-molecule events were identified using an all-photon-burst-search algorithm with a threshold of 15, a time window of 500 μs and a minimum total photon number of 150 ^75^. E-histograms of double-labelled FRET species with Alexa555 and Alexa647 were extracted by selecting 0.25<S<0.75. E-histograms of the open state without ligand (apo) and closed state with saturation of the ligand (holo) were fitted with a Gaussian distribution 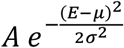.

### Visible absorption and fluorescence spectroscopy

Absorbance measurements were performed in buffer (50 mM Tris-HCl pH 7.4, 50 mM KCl) on a continuous-wave UV/VIS spectrometer (LAMBDA 465, Perkin Elmer). Absorbance spectra were recorded at a maximum absorbance of ~0.4 and base-line corrected to remove background.

Fluorescence emission was recorded in buffer (50 mM Tris-HCl pH 7.4, 50 mM KCl) on a fluorescence spectrometer (LS 55, Perkin Elmer) with excitation/emission slit width of 5 nm and gain values set to 775 V (PMT R928, Hamamatsu). The spectra were corrected for wavelength-dependent detection efficiencies.

For data representation and Förster radius calculation, the mean of three repeats of absorbance and emission spectra was calculated and normalized.

### Quantum yield measurements

For quantum yield measurements, three dilution series at 5 different concentrations were recorded in absorbance and emission. The absorbance value at the excitation wavelength was averaged over the interval 510±2.5 nm for green and 610±2.5 nm for red fluorophores. The integrated fluorescence was calculated according to 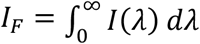. The respective absorbance values *A_λ_ex__* at 510 nm (green fluorophore) and 610 nm (red fluorophore) were fitted to the function 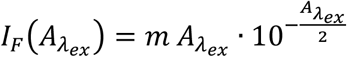, where the factor 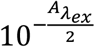 accounts for the absorption of the excitation light of the emission spectra measurements.

The fluorescence quantum yield of the fluorophores is calculated from the slopes of the fits *m_ref_* and a reference fluorophore *m_ref_* as

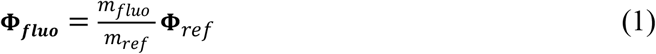

where **Φ**_*ref*_ = 91% was Rhodamine 6G^76^ for the green fluorophores and **Φ**_*ref*_ = 33% was Alexa Fluor 647 as reference^77^ for red dyes. The reported values and standard deviations result from three independent experiments.

### Förster radius calculation

The Förster radius *R*_0_ was calculated according to

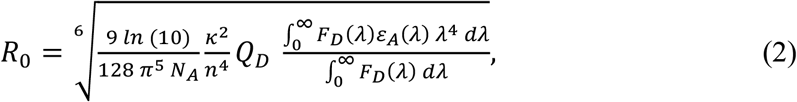

with the following values set to theoretical values:

Orientation factor *κ*^2^: 2/3
Averaged refractive index n: 1.33 in buffer / 1.4 for protein (ref. ^78^)
Extinction coefficient at maximum *ε_A_max__*: 270 000 1/(M cm) (ref. ^77,79^)

The other parameters were derived from absorption/emission spectra and quantum yield measurements as described above.

### Time-correlated single-photon counting for lifetime and anisotropy determination

Bulk lifetime and polarization decay measurements were performed using on a homebuilt setup (Supplementary Figure 3a) as also described in ref. ^80^ (Chapter 11): 400 μl of sample was measured in a 1.5×10 mm cuvette at a concentration of around 100 nM. The samples were excited by a pulsed laser (LDH-P-FA-530B for green fluorophores/LDH-D-C-640 for red fluorophores with PDL 828 “Sepia II” controller, Picoquant, GER). Excitation polarization was set with a lambda-half-waveplate (ACWP-450-650-10-2-R12 AR/AR, Laser Components) and a linear polarizer (glass polarizer #54-926, Edmund Optics). Emission light was polarization filtered (wire grid polarizer #34-315, Edmund Optics). The emission light was collected with a lens (AC254-100-A, Thorlabs) and scattering light or Raman contributions were blocked with filters (green: 532 LP Edge Basic & 596/83 BrightLine HC, AHF; red: 635 LP Edge Basic & 685/80 ET Bandpass, AHF). The signal was recorded with an avalanche-photo-diode (SPCM-AQRH-34, Excelitas) and a TCSPC module (HydraHarp400, Picoquant). Polarization optics were mounted in homebuilt, 3D-printed rotation mounts and the APD is protected from light with a 3D-printed shutter unit. An additional neutral density filter with OD = 4 in combination with a flip-mirror was used to guide the laser directly into the detection path for the measurement of the instrument response function.

In a typical experiment, the excitation power was set to 10 μW at a repetition rate of 20 MHz. The sample concentration was always tuned to obtain a ~50 kHz photon count rate. For anisotropy and lifetime measurements, data sets were recorded for each polarization setting for 5 min in the order vertical (VV1), horizontal (VH1), magic angle (MA), horizontal (VH2), and vertical polarization (VV2) under vertical excitation. The anisotropy was calculated based on the sum of two vertical and horizontal measurements to compensate for small drifts in laser power or slow changes in fluorophore concentration due to sticking. With *VV*(*t*) = *VV*1(*t*) + *VV*2(*t*) and *VH*(*t*) = *VH*1(*t*) + *VH*2(*t*), we obtained the anisotropy decay as 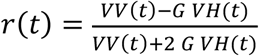, where *G* is the correction factor obtained by measuring with horizontal excitation *G* = *HV/HH* (*HV* and *HH* is the total signal in the vertical or horizontal channel, respectively)^80^.

The IRF was approximated as a sum of (up to) 3 Gaussians convoluted with a fast exponential decay, which fitted and reproduced the IRF in our setup:

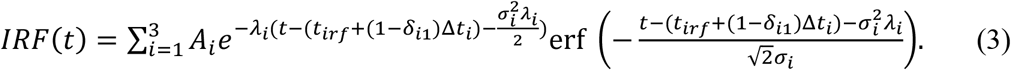

The times of the Gaussian-exponential convolutes for *i* > 1 are defined as relative time shifts Δ*t_i_* with respect to *t_irf_* in order to enable that the instrument response function can be shifted with one single time parameter *t_irf_*. Please note that the choice for this function was due to the possibility to analytically convolute the IRF with exponential decays. Alternatively, other functions could be used to describe the IRF, e.g., a sum of gamma distribution, with the same benefit. Alternatively, well-established numerical re-convolution fits could have been performed^80,81^. For our system, however, the fits were more robust with respect to small IRF mismatches with the described analytical approach.

The parameters were derived from a fit of (*IRF*(*t*) + *bkg*) to the measured instrument response function (Supplementary Figure 3b). The lifetime decays were fitted as convolution of the background-free IRF with a single (N = 1) or double exponential decay (N = 2), were the fitted IRF parameter were fixed, except of *t_irf_* to compensate for small shifts due to heating/cooling effects (Supplementary Figure 3c/e).

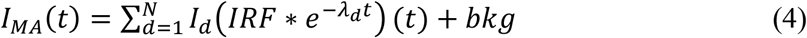

The polarization intensities read as

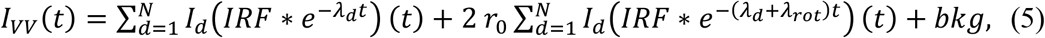

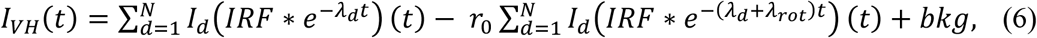

(see also ref. ^82,83^) where the parameters *I_d_, λ_d_, t_irf_*, and *bkg* are fixed. The calculated anisotropy was fitted with the model 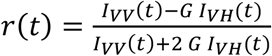, where the inverse rotational correlation time *λ_rot_* and the intrinsic anisotropy *r*_0_ are the only free parameter (Supplementary Figure 3d/f). All fits were performed as least square fits with weighted residuals according to Poissonian photon statistics.

### Mass spectrometry

For mass spectrometric analysis fluorophore standards were run on an ultra-high-performance liquid chromatographic (UHPLC) system including a diode array detector array detector (DAD) (Dionex Ultimate 3000 UHPLC, Thermo Fisher Scientific, Waltham, USA) coupled to a timsTOF MS (Bruker Daltonik, Bremen, Germany). Five microliter of each fluorophore sample was injected and separated using a C8 reversed phase column (Ultra C8, 3 μm, 2.1 x 100 mm, Restek GmbH, Bad Homburg, Germany) with 300 μl flow per minute at 60°C. Solvents were water (A) and a mixture (70/30 v/v) of acetonitrile and isopropanol (B), both containing 1% ammonium acetate and 0.1% acetic acid. The gradient started with 1 min at 55% B followed by a slow ramp to 99% B and a fast ramp within 14 min. This was kept constant for 7 min and returned to 55% B with additional 4 min of re-equilibration.

Using the DAD the absorption spectra of 190-800 nm were recorded. In parallel mass spectra were acquired by otofControl 4.0 in negative MSMS mode from 100-1300 m/z mass range. The most important parameters are set as followed: capillary voltage 4000 V, nebulizer pressure 1.8 bar, nitrogen dry gas 8 l min-1 at 200°C, collision energy 70 eV, Collision RD 800 Vpp (volt peak to peak). The evaluation was performed by Data Analysis 4.5and MetaboScape 4.0. All software tools were provided by Bruker (Bruker Daltonik, Bremen, Germany).

### Molecular dynamics simulations

Molecular dynamics (MD) simulations of mutants A186C and S352C of the *E. coli* maltose binding protein (PDB ID 1OMP^84^), each labelled with either of the three fluorophore structures AF555, Alexa Fluor 555 (structural variant #1)) and Alexa Fluor 555 (structural variant #2) at the mutated site, were performed using the GROMACS MD simulation engine^85^. The initial structures of the fluorophore-labelled proteins were built in PyMOL^86^. The fluorophore was initially oriented away from the protein. For the protein, the amber99sb^87^ force-field description was used. Fluorophore parameters were obtained as follows. The fluorophore was cut off including the linking cysteine residue and the cysteine termini were capped with an N-methyl amide group at the C-terminus and an acetyl group at the N-terminus. The AM1 method^88^in the AMBER antechamber package^89^ was used to optimize the geometries and determine partial charges for the three dye structures, as well as to determine atom types based on the gaff force field^90^. The resulting partial charges are very similar in equivalent functional groups in the three dyes (Supplementary Figure SM1). The partial charges assigned to the sulphonated groups SO_3_^-^ were found to be similar to other existing dye parameterizations^91^ (Supplementary Table 1, Supplementary Figure S4/5). If available, covalent interaction terms were taken from the AMBER-DYES force field^60^. Missing terms were taken from the gaff-based antechamber parameterization.

The fluorophore-labelled proteins were solvated in cubic computational water boxes of edge length 9.5-10.2 nm. The TIP3P water model^92^ was used. A neutralizing amount of sodium counterions was added to the solvent. The systems were energy-minimized using the steepest descent algorithm. Position restraints with a force constant of 1000 kJ mol nm^-2^ were put on all solute heavy atoms and the system was simulated for 100 ps at constant volume and a temperature of 100 K. Throughout, the Berendsen thermostat^93^ with a coupling time of 0.1 ps was used for temperature control. In a second and third equilibration step, the system was simulated with reduced (force constant 500 kJ mol nm^-2^) and vanishing position restraints, respectively, at temperatures of 200 and 300 K, respectively, for 100 ps. In a final equilibration step of 100 ps length, pressure control was introduced via the Berendsen barostat^93^ using a target pressure of 1 bar, a coupling time of 1.0 ps and an isothermal compressibility of 4.5·10^-5^ bar^-1^. For all MD simulations, a time step of 0.002 ps was used, bond lengths were kept constant with the LINCS algorithm^94^, van der Waals interactions were described with the Lennard-Jones potential^95^ and a cutoff of 1.4 nm and electrostatic interactions were described with the reaction-field method^96^, a cutoff of 1.4 nm and a dielectric constant of 80. Coordinates were written to file every 6 ps.

For each of the six equilibrated systems, four long production runs at 300 K and 1 bar, differing in the set of initial velocities assigned from the Maxwell-Boltzmann distribution, of 200 ns length were performed. From these simulations, the minimum distances between the SO_3_^-^ sulfur atoms and any protein heavy atom were determined. Distinct fluorophore-dependent behavior concerning the terminal SO_3_^-^ in the indole ring attached to the protein linker was detected which is why a set of 19-22 configurations were sampled from the compiled 800 ns simulations per fluorophore-protein system such that these configurations reflect the 800 ns-simulation data in terms of the probability distribution of minimum distances between the terminal SO_3_^-^ in the indole ring attached to the protein linker and any protein heavy atom (Supplementary Figure S6). These structures were used as initial structures in multiple short simulations (20 ns) to calculate the rotational anisotropy decay,

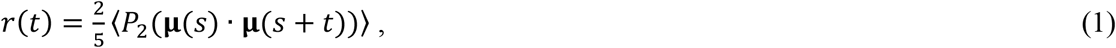

where *P*_2_(*x*)=(3*x*^2^-1)/2 is the second Legendre polynomial and **μ**(*t*) is the transition dipole moment vector at time *t* and the averaging denoted by angular brackets is done over time origins^97,98^.

## 3. Results

### Spectroscopic characterization of Alexa and AF dyes

We started our investigation of Alexa Fluor 555 and Alexa Fluor 647 properties by a comparison of absorbance and fluorescence spectra and the determination of spectroscopic parameters such as fluorescence lifetime and anisotropy against well-characterized green dyes such as Cy3, sulfo-Cy3, AF555 (cyanines), Alexa546 (rhodamine). For comparison of Alexa Fluor 647, we selected Cy5, sulfo-Cy5, AF647 (cyanines) and ATTO647N (carbopyronine); data see Figure 2.

**Figure 2.**
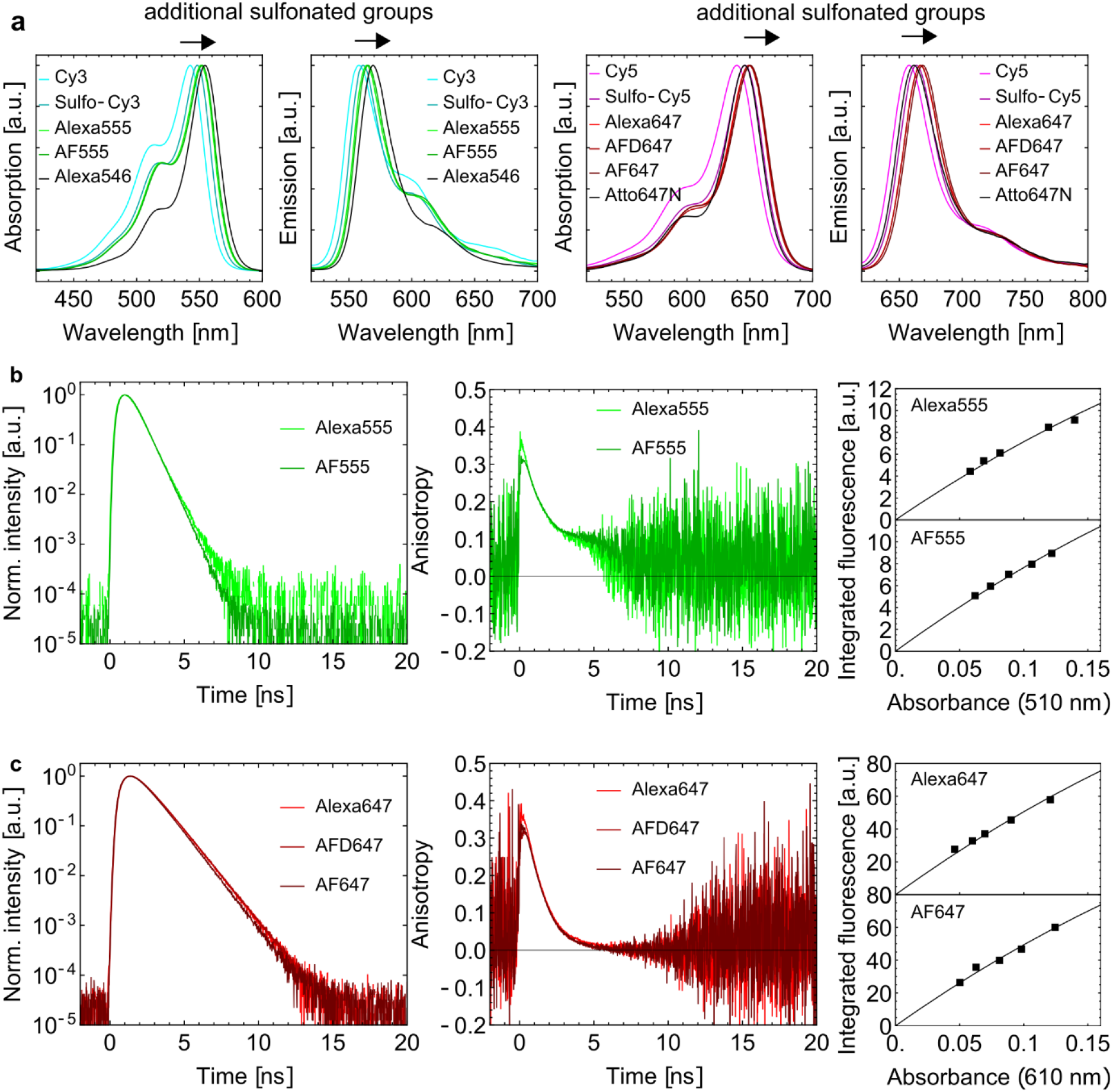
Spectroscopic characterization of bulk solutions of free green and red cyanine fluorophores. **(a)** Absorbance and emission spectra of Alexa Fluor 555 and AF555 in comparison to Cy3, Sulfo-Cy3 and Alexa Fluor 546 (left) show red-shifted spectra for increased number of SO_3_^-^-groups and difference in spectral shape compared to the rhodamine-derivative Alexa Fluor 546. Absorbance and emission spectra of Alexa Fluor 647, AFD647 and AF647 in comparison to Cy5, Sulfo-Cy5 and Atto647N (left) show red-shifted spectra for increased number of SO_3_^-^groups with small difference in spectral shape compared to Atto647N. **(b)** Lifetime (left) and time-resolved anisotropy measurements (right) of free Fluor Alexa 555 (lighter green) and AF555 (darker green) at 100 nM concentration. **(c)** Lifetime (left) and time-resolved anisotropy measurements (right) of free Fluor Alexa 647 (lighter red) and AFD647 (darker red) at 100 nM concentration. **(d)** Integrated intensity versus absorbance at 510 nm at five different concentrations (squares) for Alexa Fluor 555 (top) and AF555 (bottom) with absorbance-corrected curve fit (solid line, see methods). **(e)** Integrated intensity versus absorbance at 610 nm at five different concentrations (squares) for Alexa Fluor 647 (top) and AF647 (bottom) with absorbance-corrected curve fit as in (c).

By inspection of normalized spectra of the green-absorbing dyes in both absorption and emission (Figure 2a), we see a clear bathochromic shift when SO_3_^-^ groups become attached to the Cy3-core structure (Cy3→sulfo-Cy3→ AF555). All dyes show three vibronic peaks, e.g., for Cy3 at 540 nm, 510 nm and 475 nm, which are also seen for sulfo-Cy3, AF555 and Alexa Fluor 555 but at higher wavelengths. The spectra of Alexa Fluor 555 and AF555 are almost indistinguishable. These spectral characteristics of the cyanine dyes can be distinguished from e.g., rhodamine dyes such as Alexa Fluor 546 ^99^ that shows absorption and emission in a similar spectral window, but with different ratios of the vibronic levels.

Additional indication for a cyanine fluorophore-core in Alexa Fluor 555 is provided by fluorescence lifetimes experiments and relative quantum yields in comparison to AF555 (Table 1). Both the lifetime decays of Alexa Fluor 555 and AF555 and the relative quantum yields are highly similar. Any observed discrepancy was likely due to different background levels during our experiments. A reconvolution fitting procedure revealed similar lifetimes of 0.35±0.05 ns and 0.33±0.04 ns for Alexa Fluor 555 and AF555 in agreement with literature values for free Alexa Fluor 555 of 0.3 ns^29^. Time-resolved anisotropy decays also revealed comparable anisotropy decays of Alexa Fluor 555 and AF555 with steady-state anisotropies of 0.20±0.01 for both Alexa Fluor 555 and AF555. Rotational decay times of 0.40±0.04 ns and 0.45±0.04 ns were found with errors based on fit uncertainties. All this is in agreement with previously determined steady-state anisotropy values of 0.19 for Alexa Fluor 555^18^.

**Table 1:**
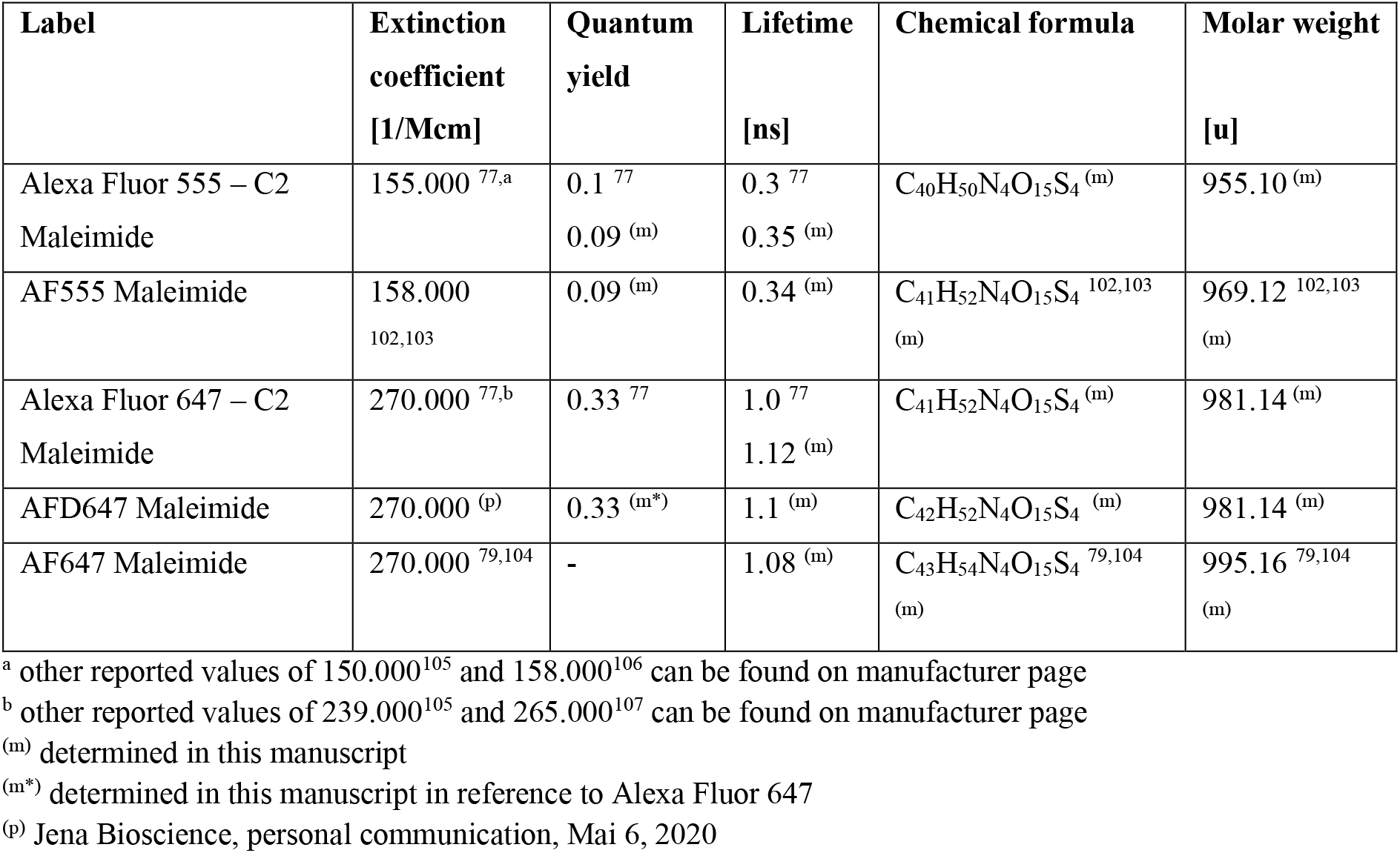
Photophysical and chemical parameters of Alexa Fluor 555/647 and AF555/AF(D)647.

| Label | Extinction coefficient<br>[1/Mcm] | Quantum yield | Lifetime<br>[ns] | Chemical formula | Molar weight<br>[u] |
| --- | --- | --- | --- | --- | --- |
| Alexa Fluor 555 – C2 Maleimide | 155.000<sup>77,a</sup> | 0.1<sup>77</sup><br>0.09<sup>(m)</sup> | 0.3<sup>77</sup><br>0.35<sup>(m)</sup> | C<sub>40</sub>H<sub>50</sub>N<sub>4</sub>O<sub>15</sub>S<sub>4</sub><sup>(m)</sup> | 955.10<sup>(m)</sup> |
| AF555 Maleimide | 158.000<sup>102,103</sup> | 0.09<sup>(m)</sup> | 0.34<sup>(m)</sup> | C<sub>41</sub>H<sub>52</sub>N<sub>4</sub>O<sub>15</sub>S<sub>4</sub><sup>102,103</sup><br>(m) | 969.12<sup>102,103</sup><br>(m) |
| Alexa Fluor 647 – C2 Maleimide | 270.000<sup>77,b</sup> | 0.33<sup>77</sup> | 1.0<sup>77</sup><br>1.12<sup>(m)</sup> | C<sub>41</sub>H<sub>52</sub>N<sub>4</sub>O<sub>15</sub>S<sub>4</sub><sup>(m)</sup> | 981.14<sup>(m)</sup> |
| AFD647 Maleimide | 270.000<sup>(p)</sup> | 0.33<sup>(m*)</sup> | 1.1<sup>(m)</sup> | C<sub>42</sub>H<sub>52</sub>N<sub>4</sub>O<sub>15</sub>S<sub>4</sub><sup>(m)</sup> | 981.14<sup>(m)</sup> |
| AF647 Maleimide | 270.000<sup>79,104</sup> | - | 1.08<sup>(m)</sup> | C<sub>43</sub>H<sub>54</sub>N<sub>4</sub>O<sub>15</sub>S<sub>4</sub><sup>79,104</sup><br>(m) | 995.16<sup>79,104</sup><br>(m) |

<sup>a</sup> other reported values of 150.000<sup>105</sup> and 158.000<sup>106</sup> can be found on manufacturer page
<sup>b</sup> other reported values of 239.000<sup>105</sup> and 265.000<sup>107</sup> can be found on manufacturer page
<sup>(m)</sup> determined in this manuscript
<sup>(m\*)</sup> determined in this manuscript in reference to Alexa Fluor 647
<sup>(p)</sup> Jena Bioscience, personal communication, Mai 6, 2020

Similar systematic trends can be observed for Alexa Fluor 647, AF(D)647 in comparison to Cy5 and Sulfo-Cy5 related to spectral shifts and variation of oscillator strength of vibronic transitions (Figure 2b). Also, the lifetime analysis of Alexa Fluor 647 and AFD647 and AF647 showed similar decays. A reconvolution fitting procedure revealed lifetimes of 1.12±0.04 ns, 1.10±0.04 ns, and 1.08±0.05 ns for Alexa Fluor 647, AFD647 and AF647, respectively, all in agreement with literature values reported for Alexa Fluor 647 of 1.0 ns^29^. Time-resolved anisotropy decays revealed comparable anisotropy decays of Alexa Fluor 647 and AFD647, and AF647 with steady-state anisotropies of 0.13±0.01 for all three fluorophores, in agreement with published values of 0.16 for Alexa Fluor 647^18^. The rotational decay time was fitted to be 0.58±0.06 ns, 0.54±0.04 ns, and 0.52±0.07 ns for Alexa Fluor 647, AFD647, and AF647, respectively. The differences in rotational correlation times were not significant for the green fluorophores (Alexa Fluor 555, AF555) and red fluorophores (Alexa Fluor 647, AFD647, and AF647).

We observed, however, a clear difference between the green fluorophores (*τ_rot_* ≈ 0.4 — 0.45 *ns*) and the red fluorophores (*τ_rot_* ≈ 0.55 *ns*), which is in good agreement with reported values for Cy3 of 0.33 ns^100^ and 0.38 ns^101^ and for Cy5 of 0.54 ns^100^. The difference can be explained by the larger size of the red fluorophores and the corresponding hydrodynamic radii (Stokes radii) of 0.75 and 0.82 nm, respectively (according to Stokes–Einstein–Debye equation under the assumption of a sphere with 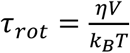, where *η* is the viscosity, *V* the sphere volume, and *k_B_T* the thermal energy). Quantum yields of Alexa Fluor 647 and AFD647 were also found to be highly similar.

Overall, our spectroscopic observations support the idea that all Alexa Fluor and AF-fluorophores studied here contain a cyanine fluorophore-core (Figure 1/2, Table 1).

### Molecular characterization of Alexa and AF dyes

To verify the molecular composition of Alexa Fluor 555 and Alexa Fluor 647, we performed mass spectrometry experiments using the AF dyes as calibration standards, since the structures of AF555 and AF647 were known. With this approach we determined the molecular mass of the malemide-derviatives of the fluorophores (Table 1, Figure 3/4) and identified characteristic molecular fragments in the MSMS spectrum based on the published structures of AF555 and AF647 (see methods). Alexa Fluor 555 maleimide showed a total mass of 955.10 u (C_40_H_50_N4O_15_S_4_), which is smaller than AF555 (969.12 u; C41H52N4O15S4) by the mass of exactly one methylene-fragment (~14 u). Also, the identified molecular fragments were very similar in terms of size and abundance when comparing Alexa Fluor 555 and AF555 with one exception: AF555 shows two fragments corresponding to sulfonated propyl- and butyl-groups, respectively. Alexa Fluor 555 does not show fragments of the sulfonated butyl-group and only a fragment related to sulfonated propyl-group(s); see Figure 3. Based on the mass spectrometry data, we were thus able to restrict the pool of potential structures for Alexa Fluor 555 to two isomers, where the maleimide linker and the sulfo-group are placed at opposing sites of the fluorophore core (see below).

**Figure 3.**
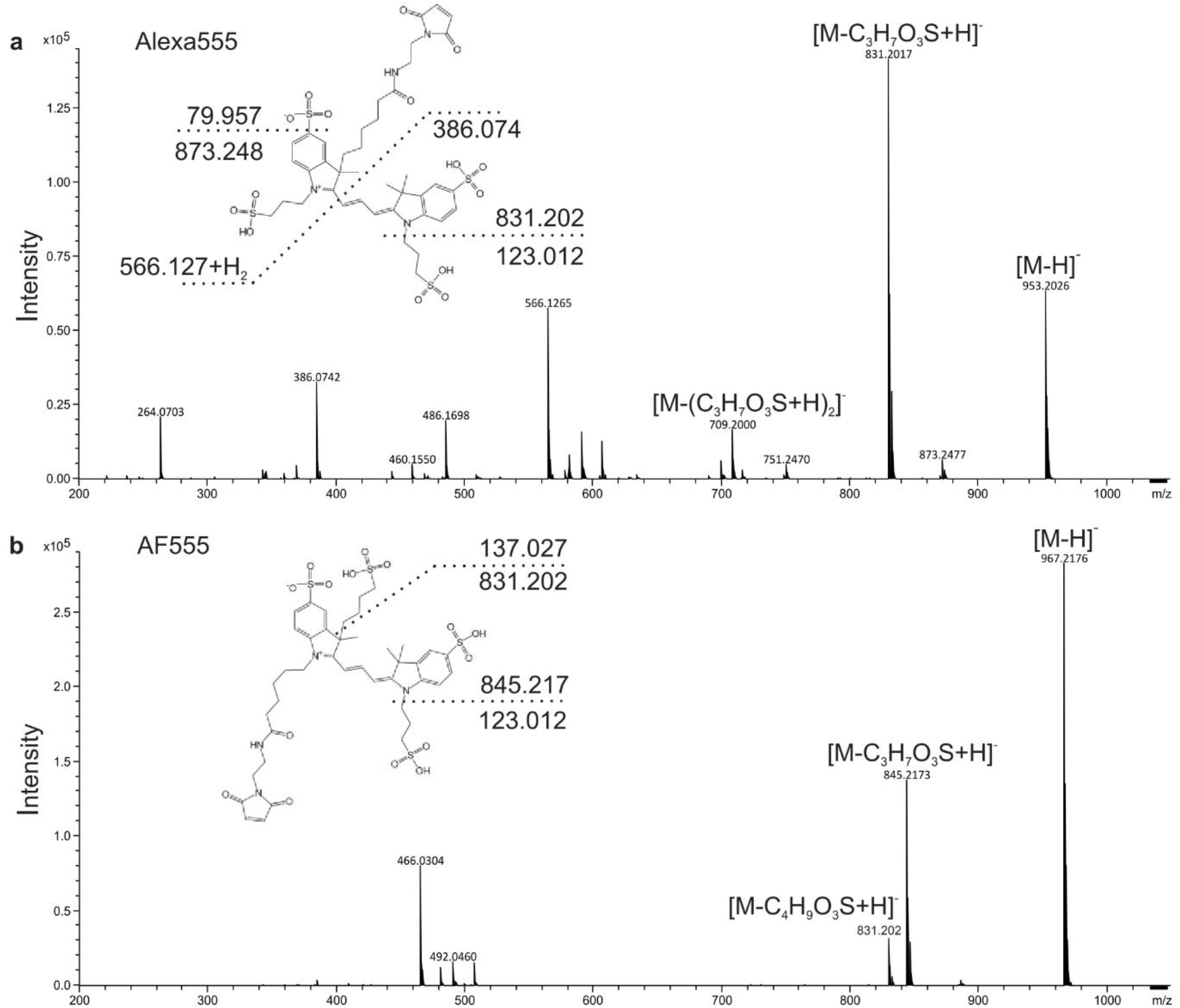
Mass spectrometry-based structure elucidation. Fragmentation mass spectra of the different fluorophores **(a)** Alexa Fluor 555 and **(b)** AF555. Mass range was set to 200-1050 m/z. Mass accuracy was 0.23 ± 0.09 ppm. Each fragmentation spectrum includes the predicted or confirmed structure. Dashed lines represent fragmentation events with resulting fragment masses or empirical formulas. M, molecular ion.

**Figure 4.**
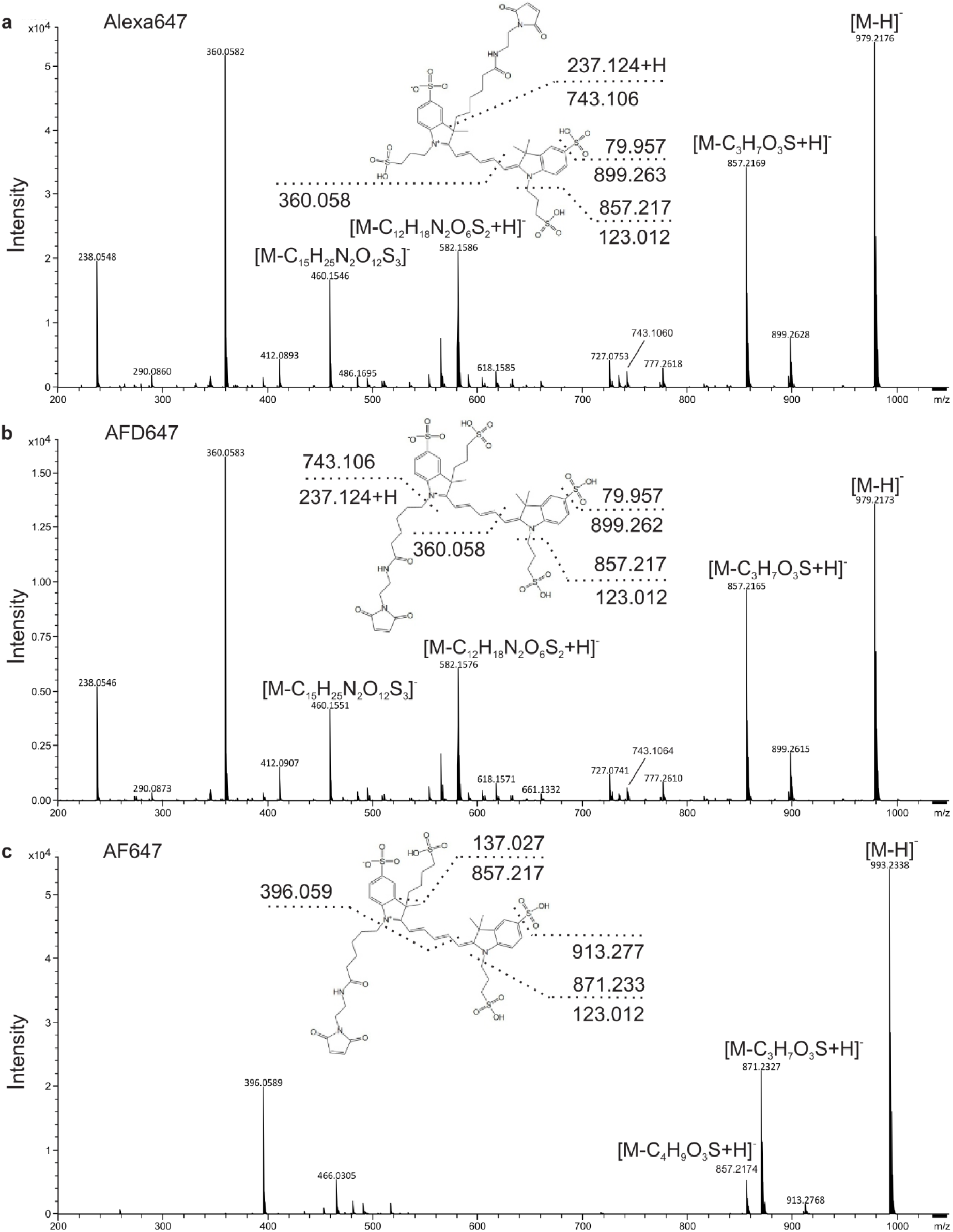
Mass spectrometry-based structure elucidation. Fragmentation mass spectra of the different fluorophores **(a)** Alexa Fluor 647, **(b)** AFD647, and **(c)** AF647. Mass range was set to 200-1050 m/z. Mass accuracy was 0.23 ± 0.09 ppm. Each fragmentation spectrum includes the predicted or confirmed structure. Dashed lines represent fragmentation events with resulting fragment masses or empirical formulas. M, molecular ion.

Using a similar approach, we compared the mass spectrometry data of AF(D)647 and Alexa Fluor 647 to verify the structure of Alexa Fluor 647 (Figure 4). The mass spectrometry data clearly shows (i.e., both total mass and fragments, Figure 4) that Alexa Fluor 647 contains two sulfonated propyl-groups attached via 3-carbon linker and the standard maleimide linker which is also used for various cyanine fluorophores including Cy3, Cy5 and its sulfonated versions (see Figure 5).

**Figure 5.**
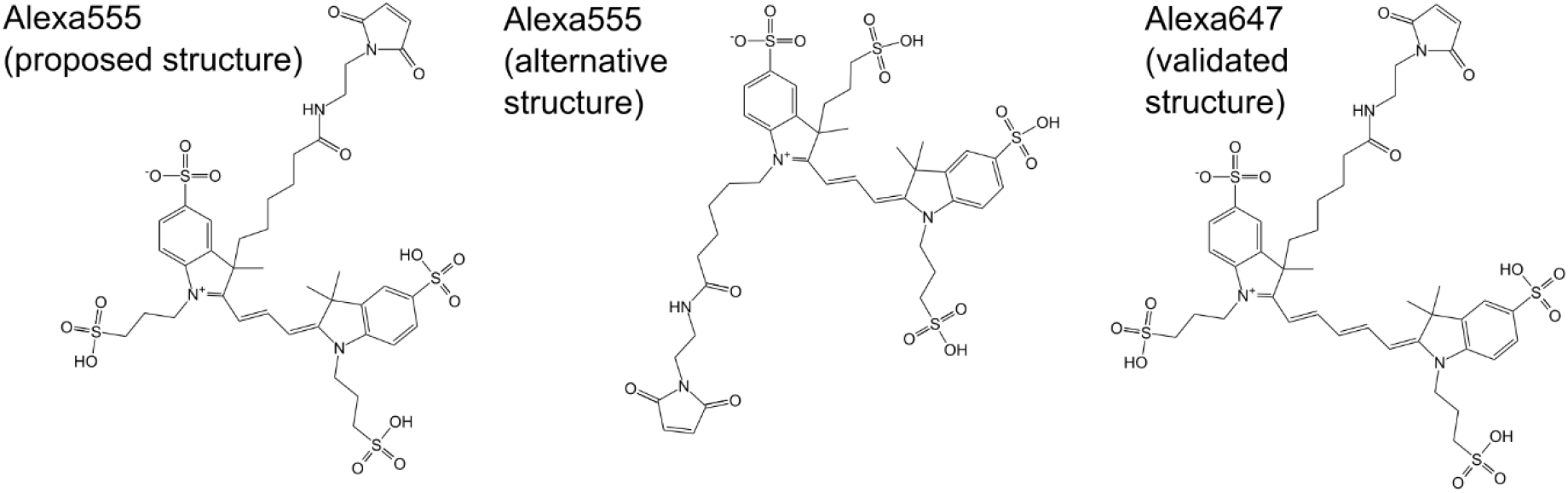
Structure determination and validation of Alexa Fluor 555 and Alexa Fluor 647. Alexa Fluor 555 has two possible structures (#1/#2). The Alexa Fluor 647 was verified including its linker structure.

Without further investigations, e.g., via NMR, both potential structures for Alexa Fluor 555 are equally probable based on our data (Figure 5). From a synthesis perspective, however, we speculate that the structural variant #1 is more likely to be correct, due to its overall similarity to Alexa Fluor 647, i.e., attachment sites of linkers and sulfonate-groups. Therefore, similar synthesis steps could be applied in production of Alexa Fluor 555 and 647. Nevertheless, we cannot completely rule out structural variant #2.

### Alexa and AF dyes for protein labelling

Next, we compared the performance of fluorophores from the Alexa Fluor and AF series for different applications in protein biophysics with the goal to use the AF dyes as a replacement of the Alexa Fluor dyes e.g., in smFRET assays. We selected the periplasmic maltose binding protein (MalE) as a model system; Figure 6a. MalE is a component of the ATP binding cassette transporter MalFGK_2_ of E. coli^108–110^. We created both single- and double-cysteine mutants of MalE (Figure 6a). These protein variants were (stochastically) labeled with fluorophores AF555, Alexa Fluor 555, AF(D)647 and Alexa Fluor 647 at strategic positions. The positions were selected to monitor ligand-induced conformational changes and related distances in MalE whenever two residues were labelled. The selected residues allowed us to create three distance pairs in MalE to monitor ligand-induced structure modulation of MalE by maltose. Two of the mutants visualized conformational motion (29-352, 87-186) and show inverse effects for addition of maltose (Figure 6b/c). We further had one MalE mutant that served as a negative control, where no ligand-induced conformational change was expected (85-352); Figure 6c. The functionality of all mutants was verified by microscale thermophoresis experiments which showed the expected ligand affinity for maltose which was in the low micromolar range (Figure S7).

**Fig. 6.**
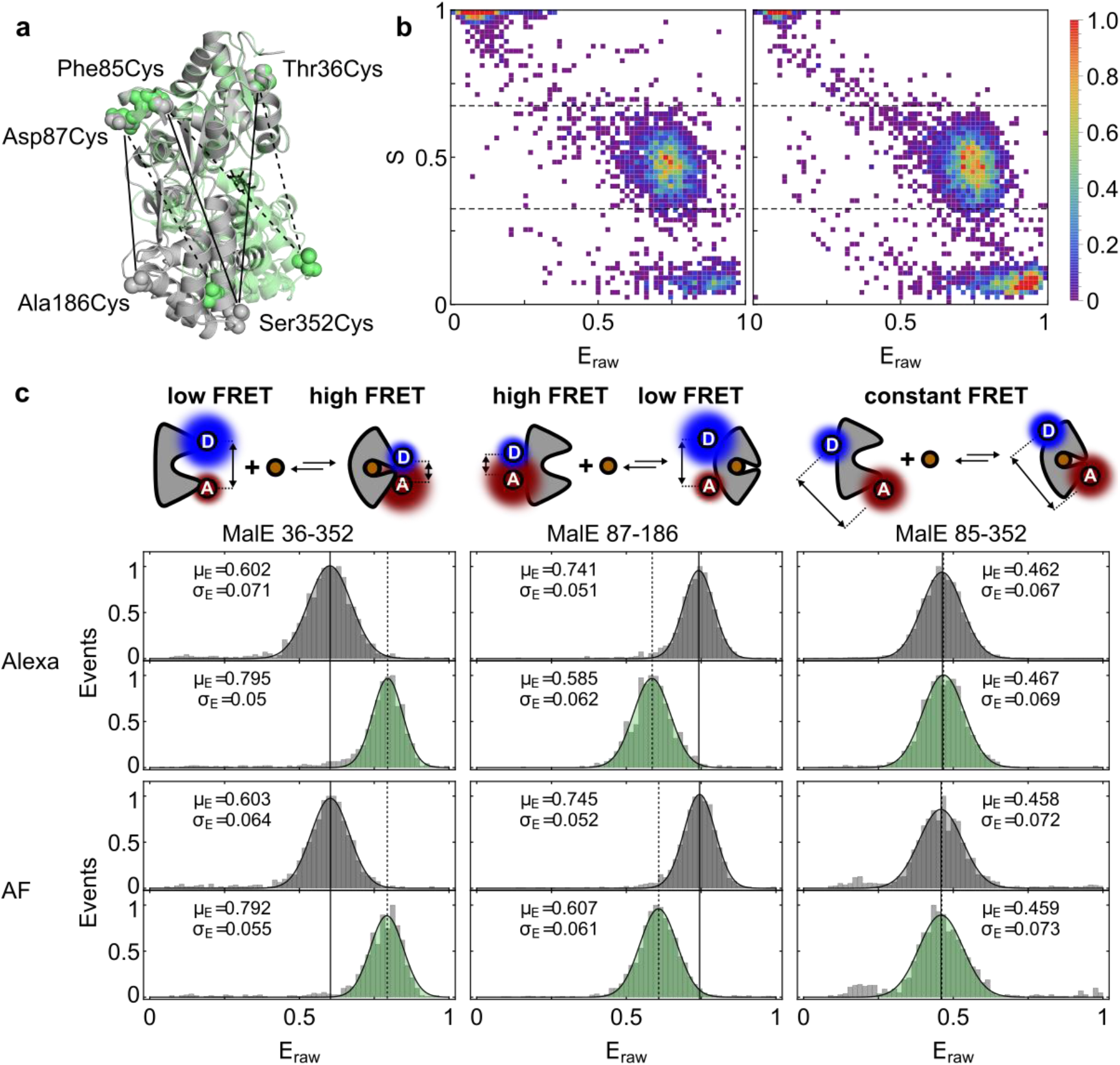
smFRET measurement comparison of Alexa Fluor 555 – Alexa Fluor 647 and AF555 – AFD647: **(a)** Overlayed crystal structures of MalE in apo state (gray, PDB 1omp) and holo state (green, PDB 1anf). Residues 4-103 are aligned and labeled residues are marked with spheres (PyMol). The FRET pairs 36-352, 87-186, and 85-352 are indicated with lines for apo (solid) and holo state (dashed). **(b)** FRET efficiency histograms (uncorrected, raw FRET values) for the three FRET mutants labeled with the Alexa Fluor pair and the AF pair are fitted with a Gaussian. The distributions show very similar FRET efficiency values for all mutants in apo state (top) and holo state with 1 mM maltose (bottom). **(c)** Representative FRET efficiency vs. stoichiometry plots (ES-plots) for MalE mutant 87-186 labeled with Alexa Fluor 555/Alexa Fluor 647 (left) and AF555/AFD647 (right) to show data quality and ratio of double labeled donor-acceptor pairs.

To define the dynamic range of FRET-assays using AF-dyes, we determined the Förster radii (R_0_) for different dye combinations based on our data. We calculated R_0_ to be 49±1 Å for Alexa Fluor 555-Alexa Fluor 647 and 50±1 Å for AF555-AFD647 in buffer with a refractive index of n = 1.33. For labeled proteins^78^, the refractive index is often assumed to an average value of n = 1.40, decreasing the Förster radii R_0_ to 47±1 Å for the Alexa- and 48±1 Å for the AF-pair. Both values are in good agreement with reported values of R_0_ for Alexa Fluor 555 and Alexa

Fluor 647 on RNA (47-48 Å)^111^ and values provided by the supplier (51 Å^77^). The distances of our selected mutants cover a substantial part of the dynamic range of smFRET for a Förster radius of ~5 nm. The cyanine nature of the dyes can, however, impose changes in the donor-quantum yield (see data in Figure 7), which implies that R0-changes variations can occur on the order of 10% provided the quantum yield does not change more than 2-fold.

**Figure 7.**
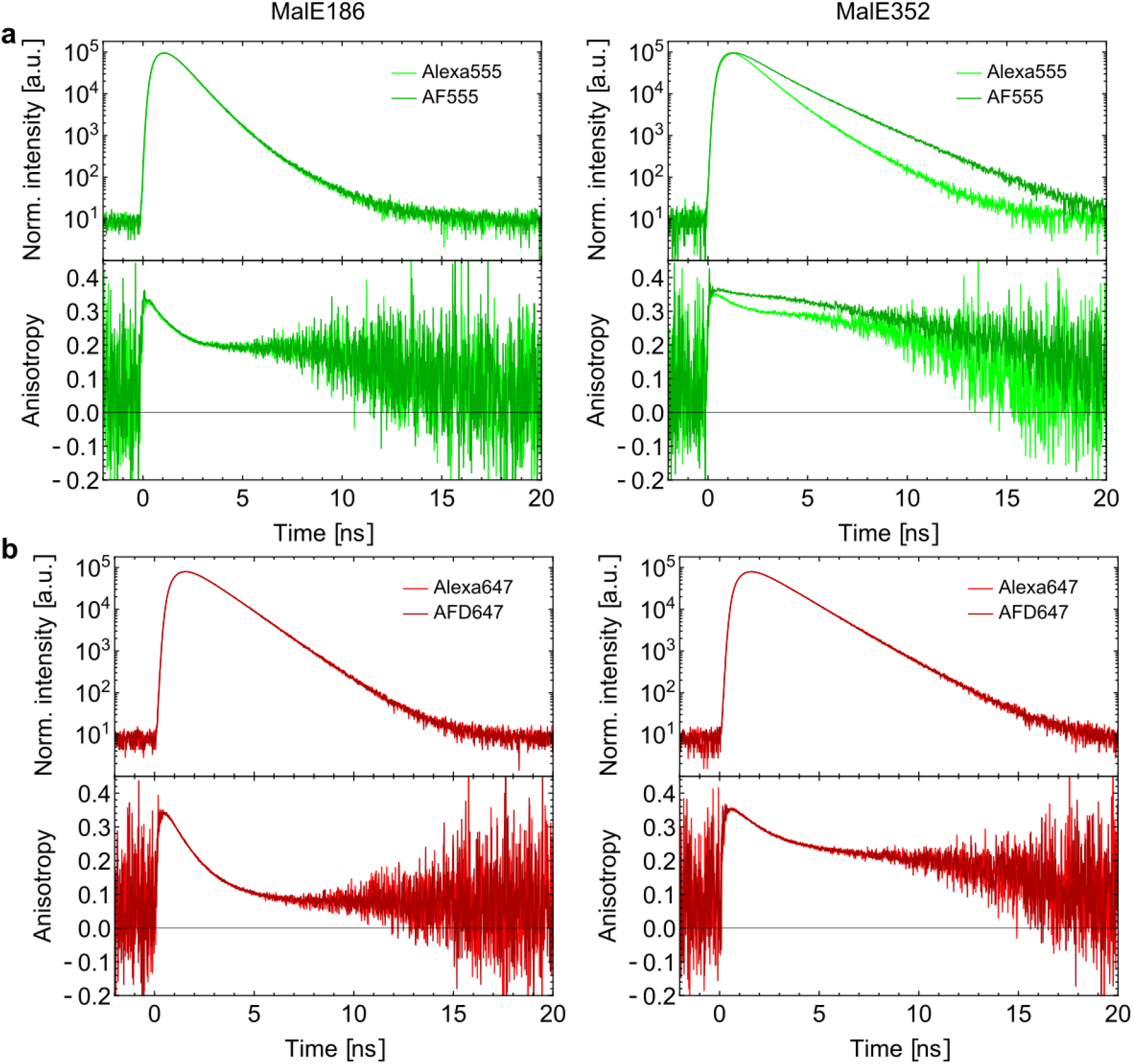
Characterization of anisotropy and lifetime decays of cyanine dyes on MalE: **(a)** Lifetime (top) and time-resolved anisotropy measurements (bottom) of Alexa Fluor Alexa 555 (lighter green) and AF555 (darker green) labelled at residues 186 (left) and 352 (right) in the ligand-free state of MalE. **(b)** Lifetime (top) and time-resolved anisotropy measurements (bottom) of Alexa Fluor Alexa 647 (lighter red) and AF647 (darker red) labelled at residues 186 (left) and 352 (right) in the ligand-free state of MalE.

### Alexa and AF dyes for smFRET studies of proteins

We benchmarked the performance of the fluorophore pair AF555-AFD647 in smFRET experiments of diffusing molecules against Alexa Fluor 555-Alexa Fluor 647 for the three different MalE variants (Figure 6). Labelling of MalE was conducted using established procedures and resulted in similar labelling efficiencies (>80%, see Supplementary Figure S1) with at least 30% donor-acceptor containing proteins. Notably, the strong interaction and sticking of AF555 at residue 352 led to a skewed profile on the size extrusion chromatogram and retarded the protein on the column (Figure S1a).

In solution-based ALEX-measurements of all three mutants, we obtained very good data quality and similar photon count rates for both dye pairs (see also Supplementary Figure S8). In a comparison of the 2D-histograms, FRET-related populations with coincident detection of donor- and acceptor-signal, we have seen only minor bleaching/blinking effects and overall shot-noise limited broadening for both dye combinations (Supplementary Figure S9/10). The generally high data quality can be seen by an inspection and comparison of the 2D-E*-S histograms, where a substantial donor-acceptor population is observed, which is also well separated from both donor- and acceptor-only species (Figure 6b). The latter is indicative of the absence of significant blinking or bleaching effects (absence of bridging populations^112^); Figure 6b.

All three double-cysteine variants also show the expected trends for the addition of ligand: low-to-high FRET (36-352), high-to-low FRET (87-186) and constant FRET (85-352); see Figure 6c. Furthermore, the mean uncorrected apparent FRET values were nearly identical for both dye pairs, i.e., their absolute E*-value varied only by about ~1%. The width of the distributions, which is characterized by the σ-values of the Gaussian fits, varied only in a moderate yet non-systematic way in between both pairs. Also, no sub-ms dynamics due to dye-photophysics were seen in burst-variance analysis (Figure S9). We noted, however, a slightly elevated bridge component for AF555/AFD647 whenever residue 352 was used as a donor dye, suggesting stronger sticking for this combination of dye/cysteine. This interpretation is further supported by the skewed SEC profile, MD simulations and time-resolved data in Figure 7 of this residue against others.

The overall high similarity of the dyes Alexa Fluor 555 / AF555 and Alexa Fluor 647 / AF(D)647 regarding their spectroscopic properties were thus faithfully reproduced in the FRET efficiency distributions that cause similar correction factors (direct excitation, leakage, quantum yield ratios) and similar Förster radii (see Figure 6). All this establishes the AF-pair as a credible alternative for smFRET investigations.

### Lifetime and anisotropy decay of Alexa and AF dyes on proteins

Notably, the smFRET experiments with the AF dyes showed a population broadening for the mutants 36-352 and 85-352 (Figure S10). To investigate this further, we characterized the environment of dyes and their protein-interactions at positions 186 and 352. We knew from previous steady-state anisotropy experiments that distinct residues in proteins can behave very differently in terms of interactions with dyes, which was also observed for MalE^52^. In general, we observed faster anisotropy decays and thus less dye-protein interactions at position 186 and stronger interactions at position 352 (slower anisotropy decay) for all four dyes (Figure 7 and Supplementary Figure S11). No detectable difference was seen for the comparison of Alexa Fluor 555 and AF555 at position 186 (Figure 7a), a result that was similar for Alexa Fluor 647 and AFD647. At residue 352, however, we identified a slow anisotropy decay indicative of strong protein-fluorophore interactions and no apparent differences between Alexa Fluor 647 and AFD647 (Figure 7b). To our surprise and despite the structural similarity of Alexa Fluor 555 and AF555, we observed significant differences in protein-fluorophore interactions between both dyes at residue 352. This is interesting since both proposed dye-structures differ mostly in the orientation of their protein-dye linker and the symmetric (Figure 5, proposed structure #1) or asymmetric placement (Figure 5, alternative structure #2) of the SO_3_^-^ residues.

Additional differences appear in the fluorescence lifetime analysis, where AF555 sticking reduces non-radiative de-excitation of the dye molecule and thus AF555 displays a longer fluorescence lifetime as compared to Alexa Fluor 555 at position 352. This effect is related to protein-induced fluorescence enhancement (PIFE)^113,114^ and supports the idea that restricted motion is responsible for higher lifetimes and increased brightness in the green cyanine fluorophores. The PIFE-effects^115,116^ observed in the comparison between Alexa Fluor 555 and AF555 provides an explanation for the PIFE caused by T7 DNA Polymerase gp5/trx with Alexa Fluor 555-labelled dsDNA (which could not be explained previously due to lacking knowledge of Alexa Fluor 555 structure). Importantly, this indicates the relevance of verifying fluorophore structures and labelling locations in relation to non-interacting environments.

To further explain the observed differences between Alexa Fluor 555 and AF555 in their interactions with MalE at specific positions, we performed molecular dynamics simulations (Figure 8). As described in the methods section, the rotational anisotropy decay *r*(*t*) was calculated for multiple short simulations of the maltose binding protein labelled with either AF555 and the two possible structures of Alexa Fluor 555 (structural variants #1 and #2).

**Fig. 8.**
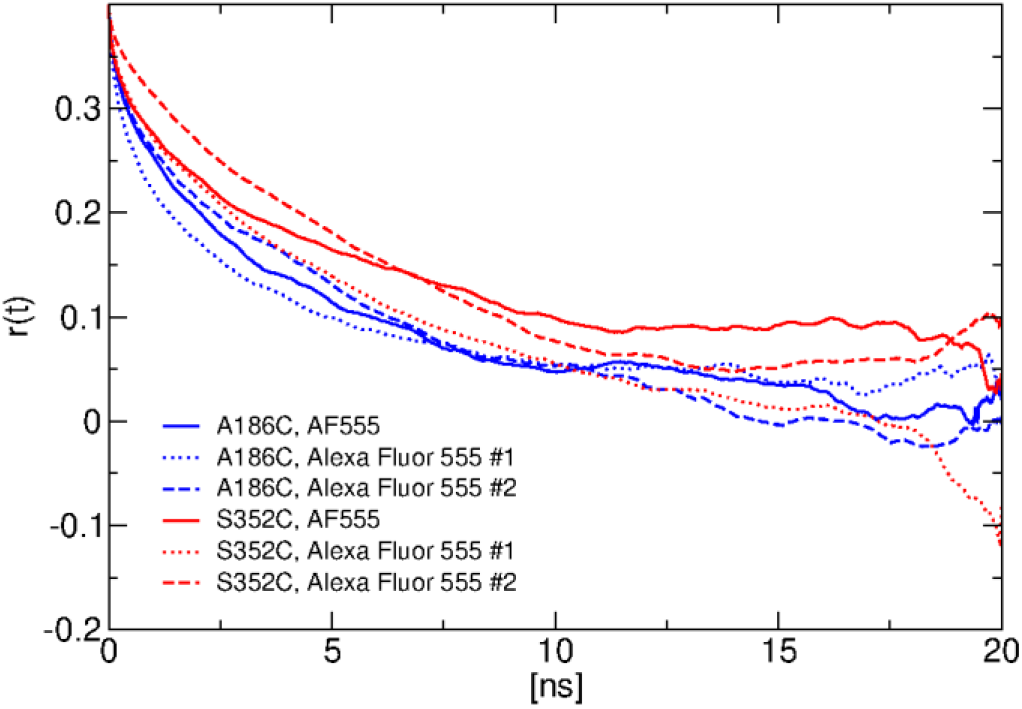
Average rotational anisotropy decay calculated from MD simulations of fluorophore-labelled MalE. Mutants A186C and S352C were combined with fluorophores AF555 and Alexa Fluor 555, the latter with the two proposed structural variants #1 and #2, were simulated for 20 ns. The curves depict the average of the anisotropy decay *r*(*t*) over ~20 simulations per protein-dye system. Time origins for the averaging per simulation are separated by 6 ps.

The rotational anisotropy decay at 186C is faster than at 352C, reflecting, as observed in experiment, reduced dye-sticking at the former site. The simulation suggests distinct interaction sites, which are very sensitive for specific structural features in the fluorophore. Important protein-dye interactions possibly contributing to the reduced motion at the 352C site and selected distances between fluorophore- and protein-atoms are illustrated in Supplementary Figure S12 and S13, respectively. Interestingly, for the 352C site, the simulation also shows a qualitative match to the experimental data for Alexa Fluor 555 (proposed structural variant #1), where a faster anisotropy decay is found (Figure 7), in significant contrast to AF555 (slow anisotropy decay; Figure 7). For Alexa Fluor 555 (proposed structural variant #2) at the 352C site, only minor differences are seen in comparison to AF555, which is not consistent with the data shown in Figure 7. The simulations thus provide further support for Alexa Fluor 555 proposed structural variant #1.

## 4. Discussion and conclusion

Using a combined investigation of the spectroscopic and molecular properties of Alexa Fluor 555, we were able to confirm that it does indeed have a cyanine fluorophore core (Figure 1–5). Similar studies on Alexa Fluor 647 allowed us to accurately determine its molecular structure and with that settle conflicting reports on its linker structure and the linkers of the sulphonated groups. Our analysis revealed two possible (isomeric) forms of Alexa Fluor 555 (Figure 5). In this submission, we only collected indirect evidence using MD simulations (Figure 8) which do not allow an unambiguous identification of the correct isomer, and a univocal assignment would require further experimental support, e.g., NMR analysis. Our spectroscopic analysis and tests of the dyes in smFRET experiments on proteins (Figure 6/7) showed good performance of all dyes in the experiments and a high degree of similarity between the Alexa Fluor and AF dyes as was expected based on the structural similarity.

Our results enabled us to derive the parameters for MD simulations (Supplementary Table 1, Supplementary Figure S4/5) and for in silico-prediction of accessible volumes of the dyes when used as a FRET label. These parameters are important for predictions of observed mean inter-fluorophore distances and FRET-efficiencies for a combination of smFRET experiments with structural modelling and simulations^117^. Using the structures of AF555 and AF(D)647 (Figure 1) and the ones of Alexa Fluor 555 and 647 (Figure 5), we derived all relevant parameters for AV calculations following the method by Kalinin et al.^117^ (Table 2). For these simulations a parametrization for linker and fluorophore core – modelled as an ellipsoid – is provided in Table 2 according to the proposed procedure (Supplementary Figure S14)^117^.

**Table 2:**
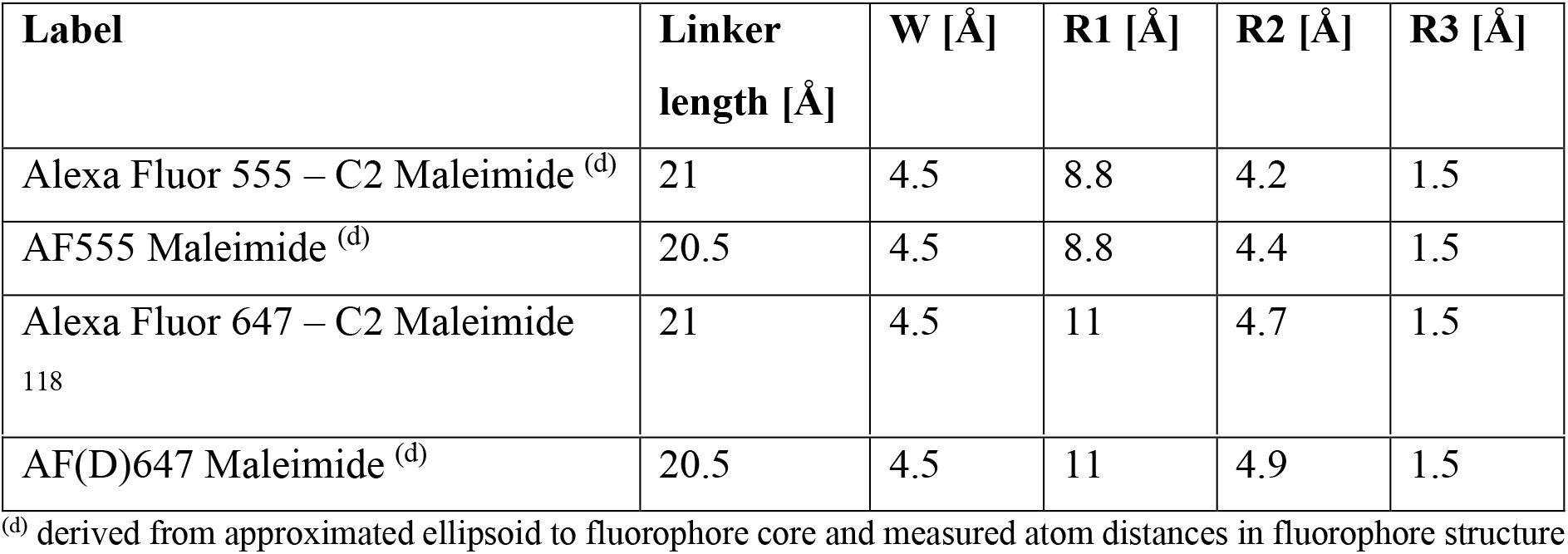
Geometric Parameters for in silico predictions of FRET labels using the FRET-restrained positioning system.

| Label | Linker length [Å] | W [Å] | R1 [Å] | R2 [Å] | R3 [Å] |
| --- | --- | --- | --- | --- | --- |
| Alexa Fluor 555 – C2 Maleimide <sup>(d)</sup> | 21 | 4.5 | 8.8 | 4.2 | 1.5 |
| AF555 Maleimide <sup>(d)</sup> | 20.5 | 4.5 | 8.8 | 4.4 | 1.5 |
| Alexa Fluor 647 – C2 Maleimide<br>118 | 21 | 4.5 | 11 | 4.7 | 1.5 |
| AF(D)647 Maleimide <sup>(d)</sup> | 20.5 | 4.5 | 11 | 4.9 | 1.5 |

<sup>(d)</sup> derived from approximated ellipsoid to fluorophore core and measured atom distances in fluorophore structure

We finally note that the high structural resemblance of the dyes might render it reasonable to use either of the dyes without considering the small differences. Yet as shown above, slight structural variations of the dyes can impact dye-protein interactions greatly, e.g., as seen for Alexa Fluor 555 and AF555 with differing lifetimes/anisotropy decays (Figure 7). Such effects, which we also observed for a comparison of Alexa Fluor 647 and AF647 (but not with AFD647) can largely alter various parameters in a biophysical assay (Figure 9). Here, we observed significant differences in lifetime and anisotropy decay for a mere addition of a methylene-bridge of the sulfonated group SO_3_^-^ (sulfonated butyl-group instead of propyl-groups). While changes in the donor lifetime can alter the Förster radius, strong dye-protein interactions can produce a large number of additional artifacts ranging from long rotational correlation times of the respective dye to an impact of the dye on the biochemical properties of the protein.

**Fig. 9.**
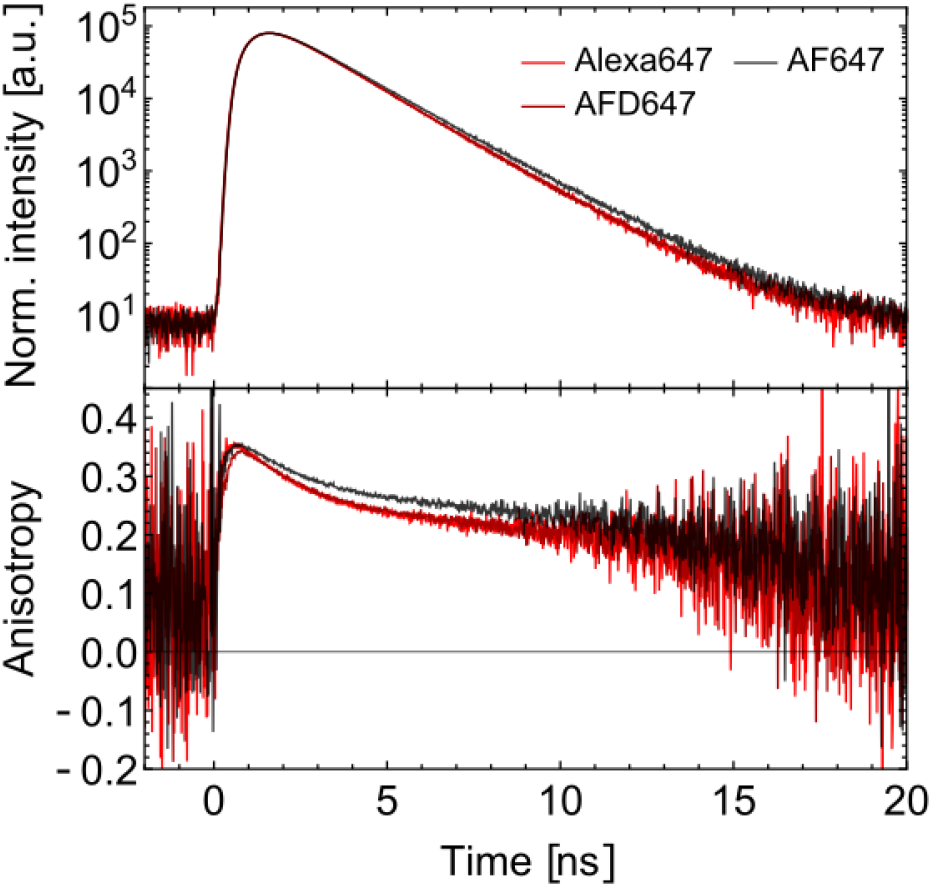
Characterization of anisotropy and lifetime decays of Alexa Fluor 647 and AF(D)647 on MalE (position 352) in the ligand-free state of MalE. The change in one CH2-group from AFD647 to AF647 shows a significant increased anisotropy and lifetime.

Overall, we conclude that the gathered structural knowledge on Alexa Fluor 555 and 647 will finally enable their applications wherever precise chemical information is required. Furthermore, we conclude that AF555 and AF(D)647 are suitable replacements of the Alexa Fluor dyes in applications for which similar spectroscopic and molecular parameters are neede.

## Acknowledgements

This work was financed by an ERC Starting Grant (ERC-StG 638536 - SM-IMPORT to T.C.), Deutsche Forschungsgemeinschaft within GRK2062 (project C03 to T.C.) and SFB863 (project A13 to T.C., project A10 and A13-111166240 to M.Z.), LMUexcellent (start-up funding to T.C.), the Center for integrated protein science Munich (CiPSM) and the Center for Nanoscience (CeNS). C.G. acknowledges a PhD fellowship from the Studienstiftung des deutschen Volkes. Compute resources for this project were partially provided by the Regionales Rechenzentrum Erlangen (RRZE).

## Competing interests

The authors declare no competing interests.

## Author contributions

C.G. and T.C. conceived and designed the study. T.C. supervised the study. C.G performed research and analyzed data. M.L. performed mass spectrometry. M.M.R. and M.Z. designed simulation setup, M.M.R performed and analyzed simulations. C.G. and T.C. wrote the manuscript. All authors discussed and interpreted the results and approved the final version of the manuscript. We finally thank D.A. Griffith for carefully reading and commenting on the manuscript.

## Supplementary Information

**Fig. S1.**
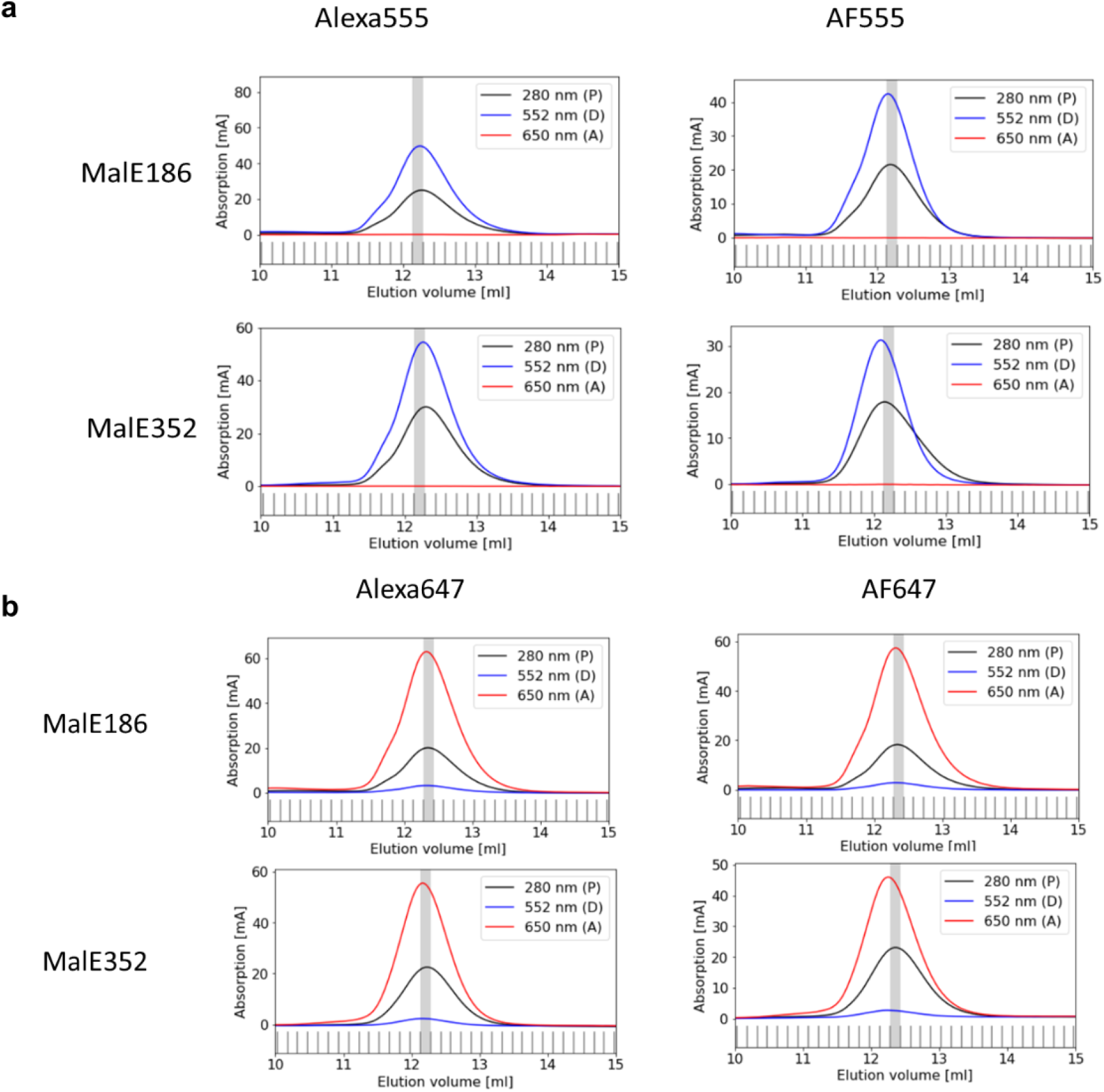
Investigation of protein quality and labeling efficiency. **(a)** Size exclusion chromatography (SEC) profiles of Alexa Fluor 555 and AF555 labeled at MalE mutants 186 and 352. **(b)** SEC profiles of Alexa Fluor 647 and AFD647 labeled at the same mutants as in (a).

**Fig. S2.**
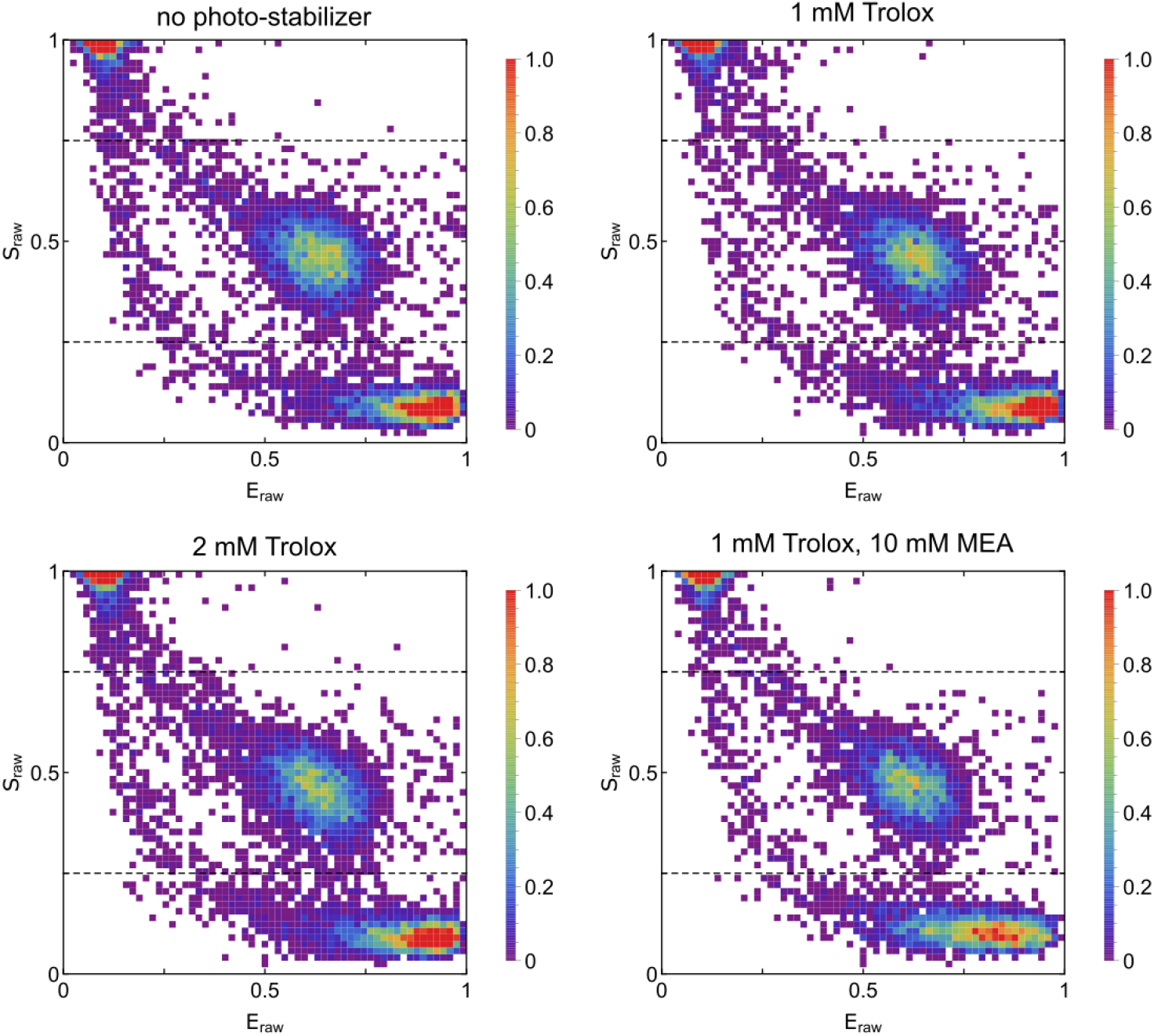
Photo-stabilizer comparison in FRET measurements with Alexa Fluor 555 – Alexa Fluor 647. ES-histogram of MalE mutant 26-352 in unbound state with photo-stabilizing agents do not show measurable differences in data quality and signal intensity.

**Fig. S3.**
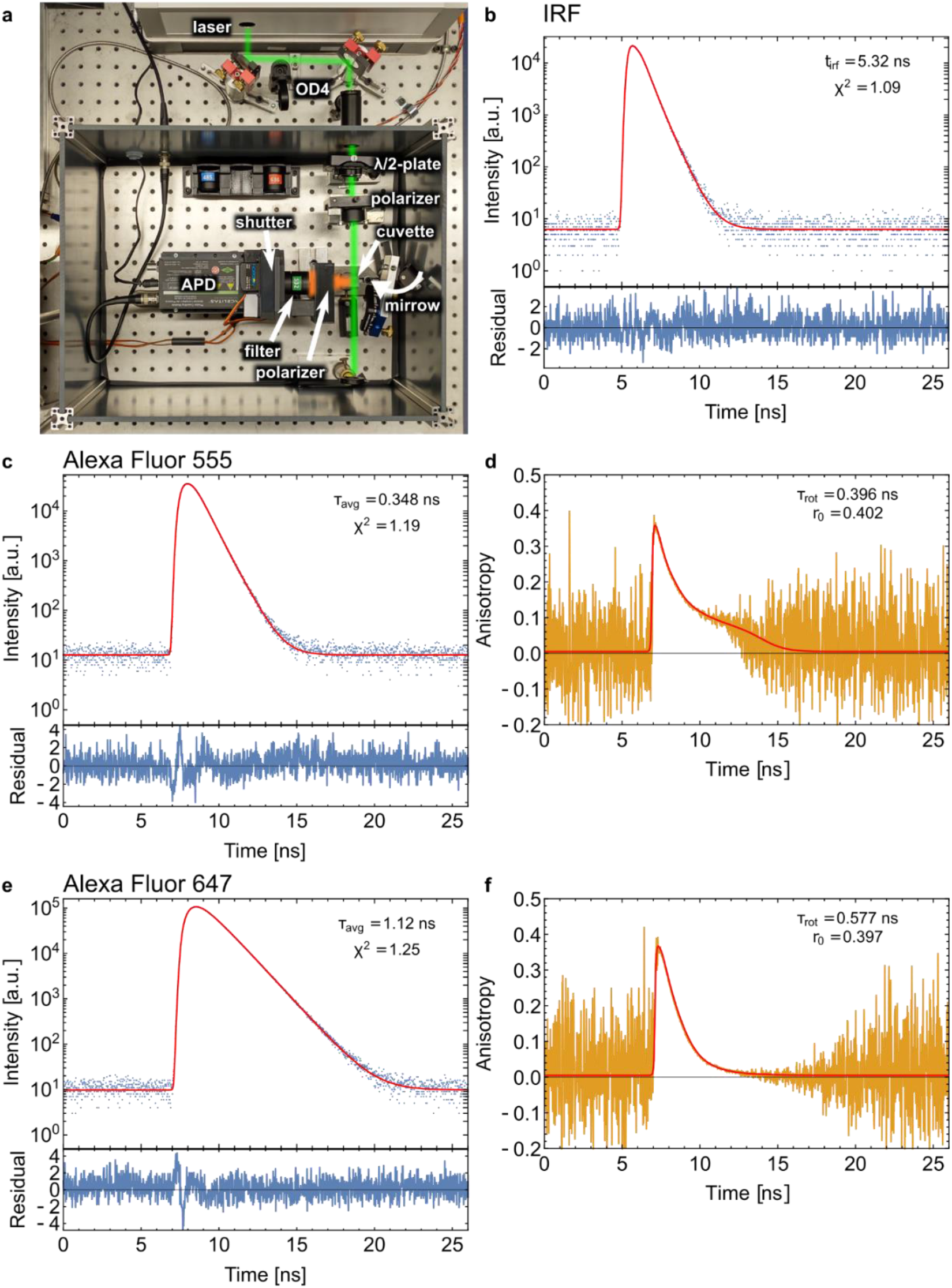
Lifetime and time-resolved anisotropy measurements: **(a)** Setup for lifetime and anisotropy measurements with labeling of the relevant parts. For IRF determination, the mirror is flipped and replaces the cuvette to guide the laser directly into the APD (with additional OD4 filter and without filter set). **(b)** Intensity profile of green laser (blue) with fitted IRF function (red) as sum of 3 Gaussians convoluted with exponential decays (top) and fit residuals (bottom). **(c)** Intensity profile of free Alexa Fluor 555 (blue) fitted with IRF function from (a) convoluted with a biexponential decay (red) (top) and fit residuals (bottom). The stated lifetime is the amplitude weighted mean of the two lifetime components. **(d)** Calculated time-resolved anisotropy of Alexa Fluor 555 (orange) fitted with reconstructed anisotropy based on intensity fit from (c) and measured detection correction factor *G* (red). The stated rotational correlation time *τ_rot_* and intrinsic anisotropy *r*_0_ are the only free fit parameters. **(e)** Intensity profile of free Alexa Fluor 647 (blue) fitted with IRF function for red laser as in (a) convoluted with a biexponential decay (red) similar to (c). **(f)** Calculated time-resolved anisotropy of Alexa Fluor 647 (orange) fitted with reconstructed anisotropy (red) similar to (d).

**Table S1:** Partial charges of the atoms of the SO_3_^-^ groups in the present fluorophore force-field description compared to the force-field descriptions of Best et al.^91^ and Graen et al.^60^. The SO_3_^-^ groups are labelled i, j depending on their position, where i = 1 denotes the indole ring attached to the protein linker, i = 2 denotes the second indole ring, j = 5 denotes a five-membered ring and j = 6 a six-membered ring. For simplicity, the partial charges are here rounded to 2 digits behind the comma. The actual unrounded charges are given in Supplementary Figure S4.

| $\text{SO}_3^-$ group | q(S)<br>[e] | q(O)<br>[e] |
| --- | --- | --- |
| AF555 |  |  |
| 1,5 | 1.42 | -0.76 |
| 1,6 | 1.46 | -0.74 |
| 2,5 | 1.44 | -0.76 |
| 2,6 | 1.46 | -0.74 |
| Alexa Fluor 555 #1 |  |  |
| 1,5 | 1.43 | -0.75 |
| 1,6 | 1.48 | -0.76 |
| 2,5 | 1.43 | -0.75 |
| 2,6 | 1.46 | -0.74 |
| Alexa Fluor 555 #2 |  |  |
| 1,5 | 1.43 | -0.76 |
| 1,6 | 1.46 | -0.75 |
| 2,5 | 1.44 | -0.76 |
| 2,6 | 1.46 | -0.74 |
| Alexa 488, Best et al. |  |  |
|  | 1.38 | -0.71 |
| Alexa 594, Best et al. |  |  |
|  | 1.46 | -0.77 |
| Alexa 350, Graen et al. |  |  |
|  | 1.09 | -0.68 |

**Fig. S4:**
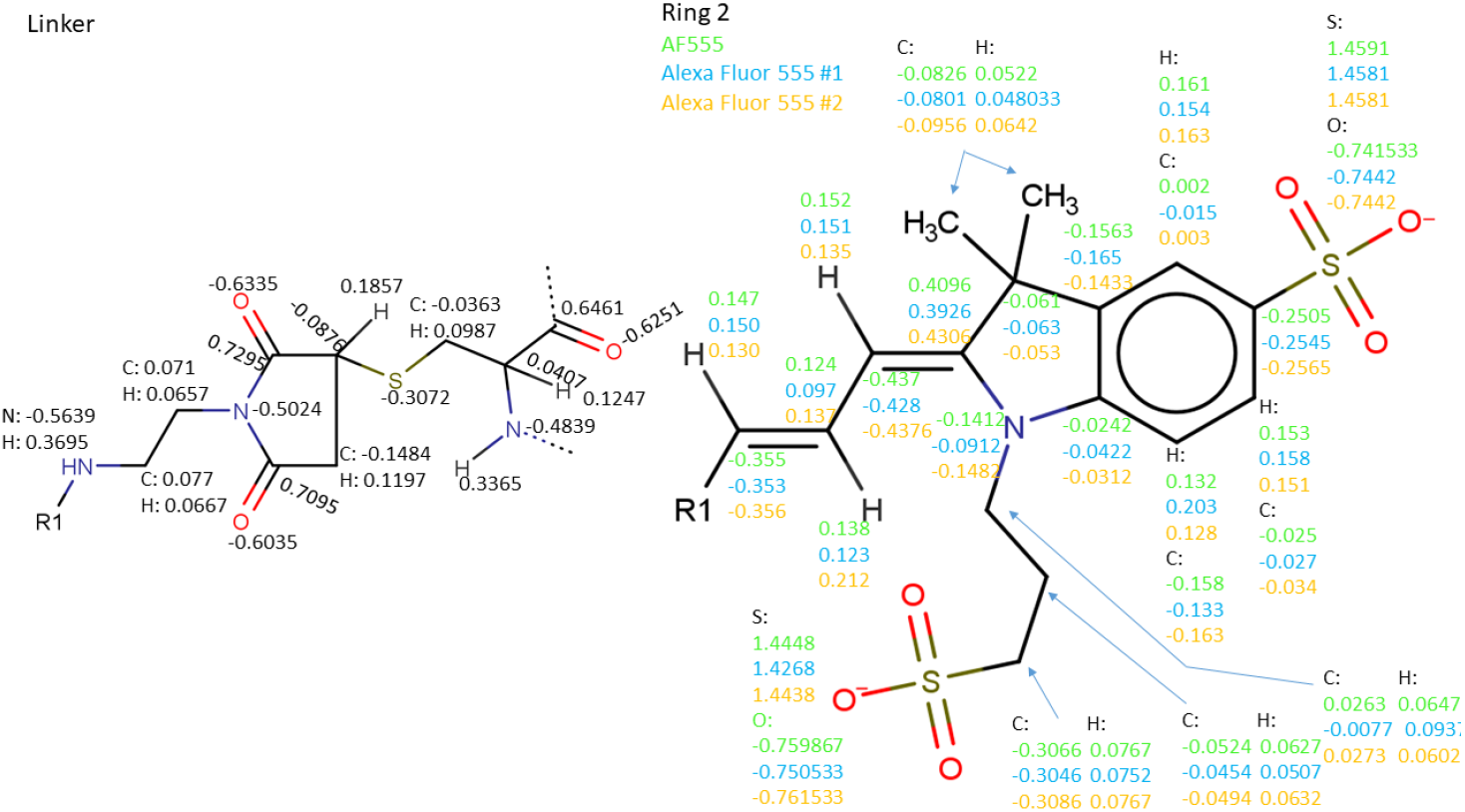
Partial charges determined for the shared structural regions of AF555, Alexa Fluor 555 #1 and Alexa Fluor 555 #2 with the antechamber package^89^ as described in the main article. Abbreviations “R”, “R1” and “R2” refer to the linker, indole ring 1 and indole ring 2 with the bridging (CH)3 groups, respectively.

**Fig. S5:**
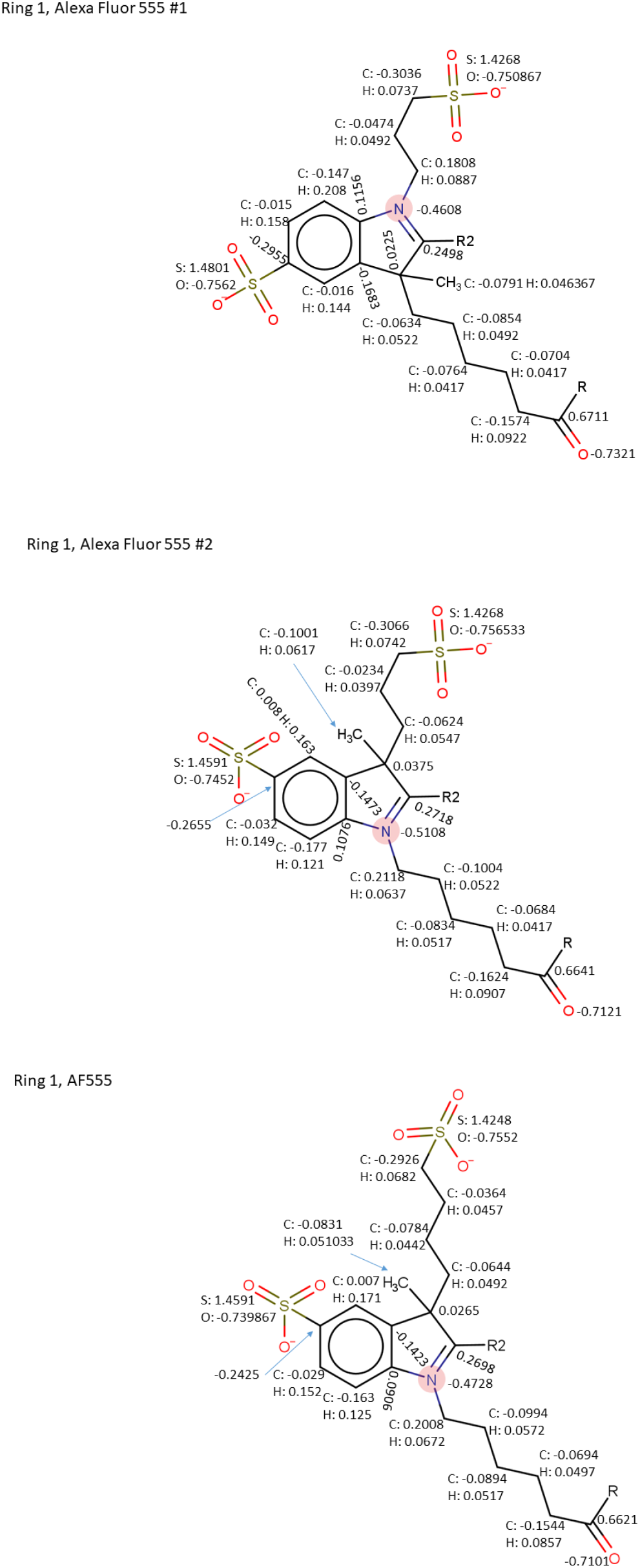
Partial charges determined for the fluorophores AF555, Alexa Fluor 555 #1 and Alexa Fluor 555 #2 in the distinct structural regions with the antechamber package^89^ as described in the main article. Abbreviations “R”, “R1” and “R2” refer to the linker, indole ring 1 and indole ring 2 with the bridging (CH)_3_ groups, respectively.

**Fig. S6:**
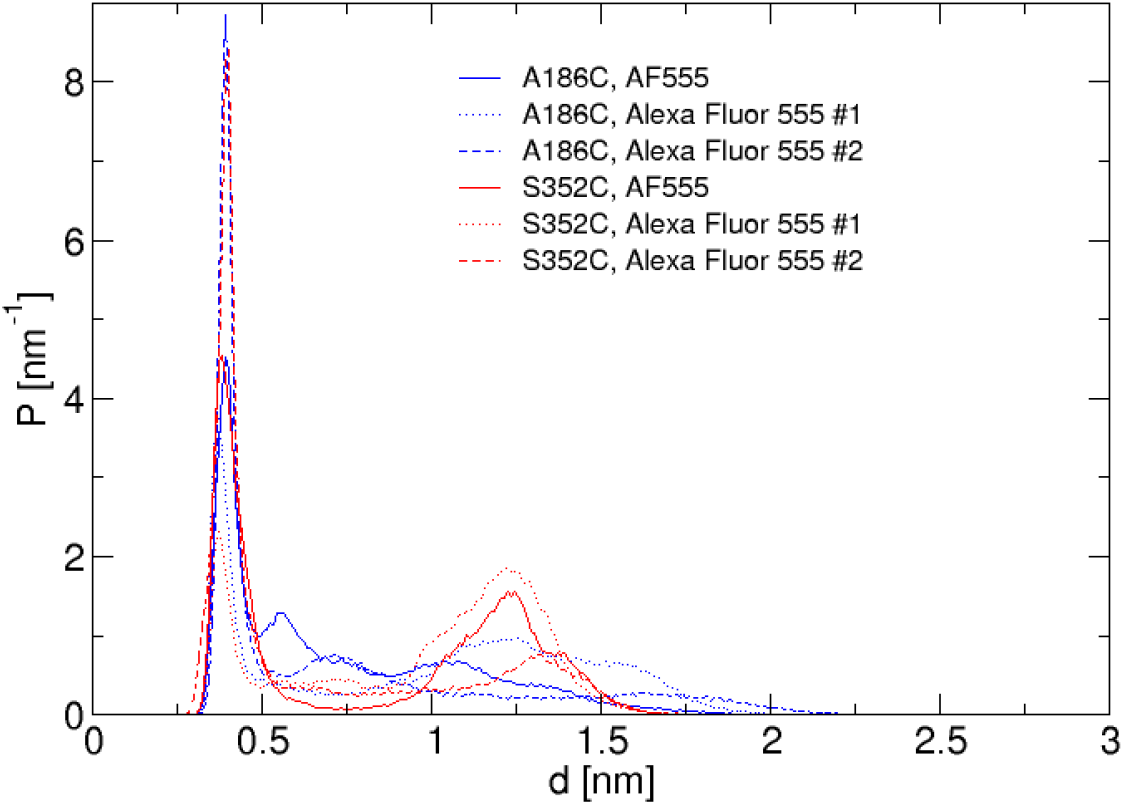
Probability distribution of the minimum distance between the sulfur atom of the terminal SO_3_^-^ group in the indole ring attached to the protein linker and any protein heavy atom, for the different dye-protein combinations, sampled during four 200 ns simulations per dye-protein combination. Large distances >1.5 nm are sampled more frequently at the A186C site than at the S352C site. At both sites, Alexa Fluor 555 #2 samples preferably small distances.

**Fig. S7.**
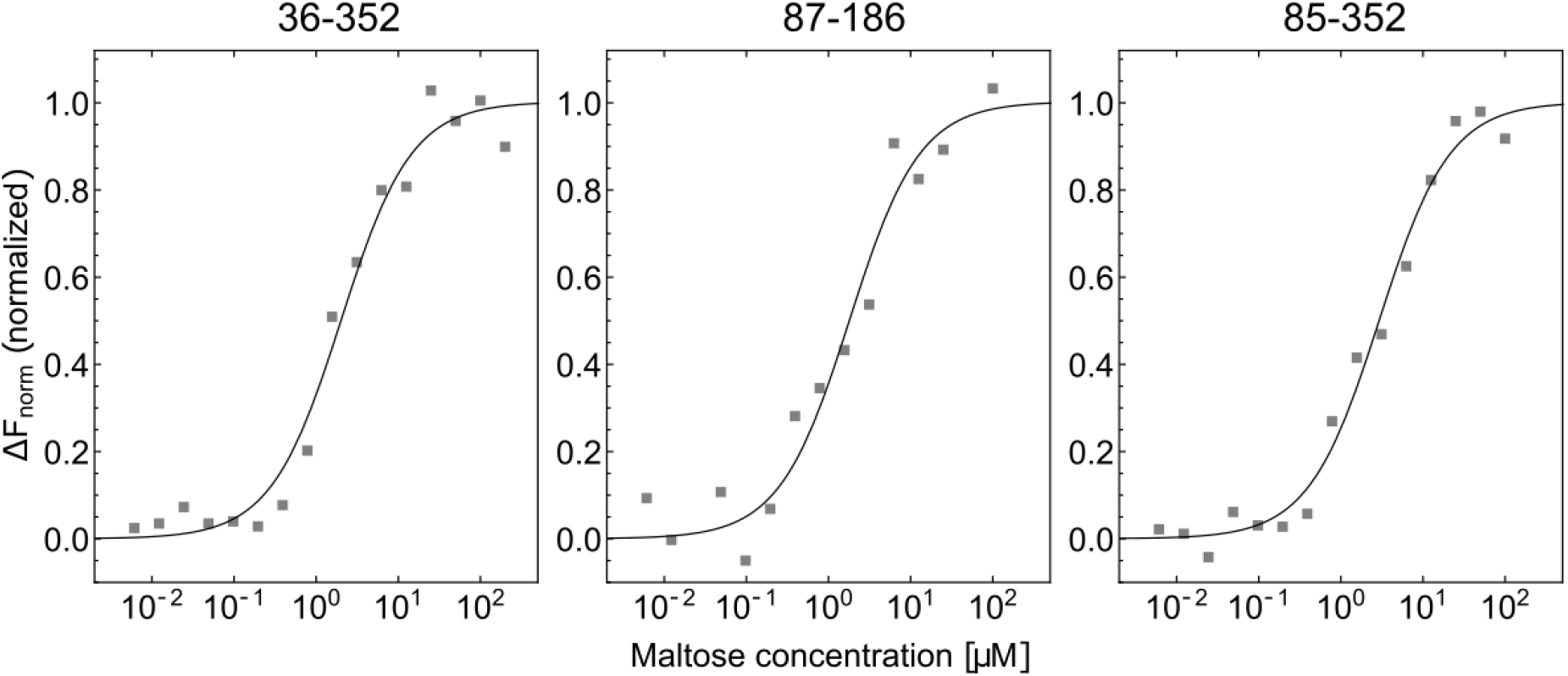
Binding affinity measurements using microscale thermophoresis. Binding affinities were measured with microscale thermophoresis (Monolith NT.LabelFree, Nanotemper), where the ratio of fluorescence before and after heating 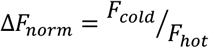 was recorded at different maltose concentrations^119^. Data points show Δ*F_norm_* normalized to minimal and maximal fluorescence intensities for unlabelled mutants 36-352 (left), 87-186 (middle), and 85-352 (right). The curves were fitted with a standard model for receptor-ligand kinetics 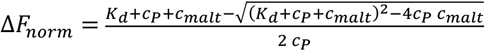, where *k_d_* is the dissociation constant, *c_P_* the protein concentration set to 0.25 μM in the experiment, and *c_malt_* the maltose concentration. The fits to the binding model (solid line) gave K_d_-values of 2.2±0.4 μM (left), 1.6±0.3 μM (middle), and 2.8±0.3 μM (right), respectively.

**Fig. S8.**
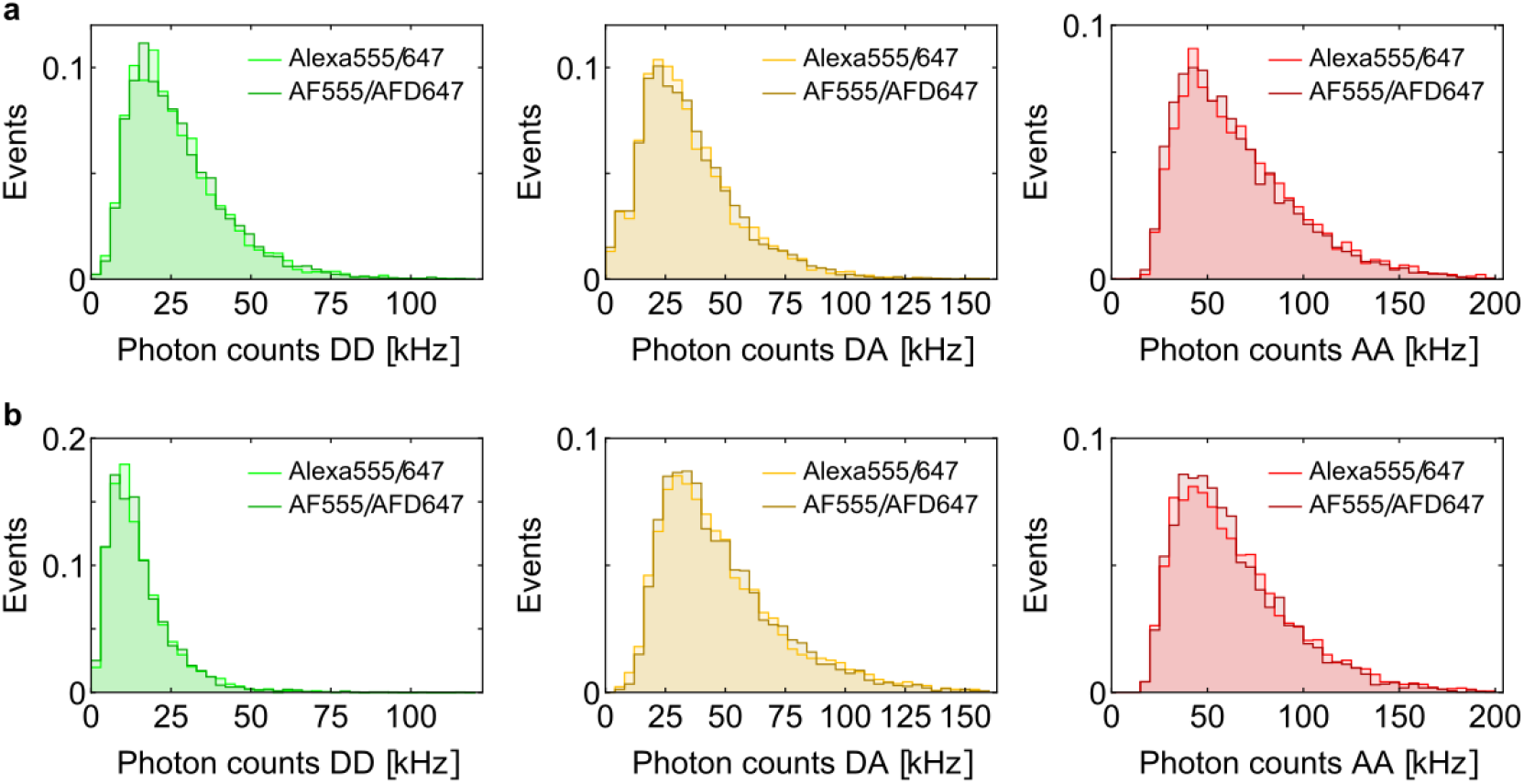
Investigation of data quality of smFRET measurements between Alexa Fluor 555/647 and AF555/AFD647: **(a)** Photon count histograms of selected FRET population (0.325<S<0.675) for MalE mutant 36-352 in apo state. The different detection channels show very similar photon rates for donor excitation – donor emission (DD, left), donor excitation – acceptor emission (DA, middle), and acceptor excitation – acceptor emission (AA, right). **(b)** Evaluation as in (a) for MalE 36-352 mutant in maltose bound state (1 mM maltose in buffer).

**Fig. S9.**
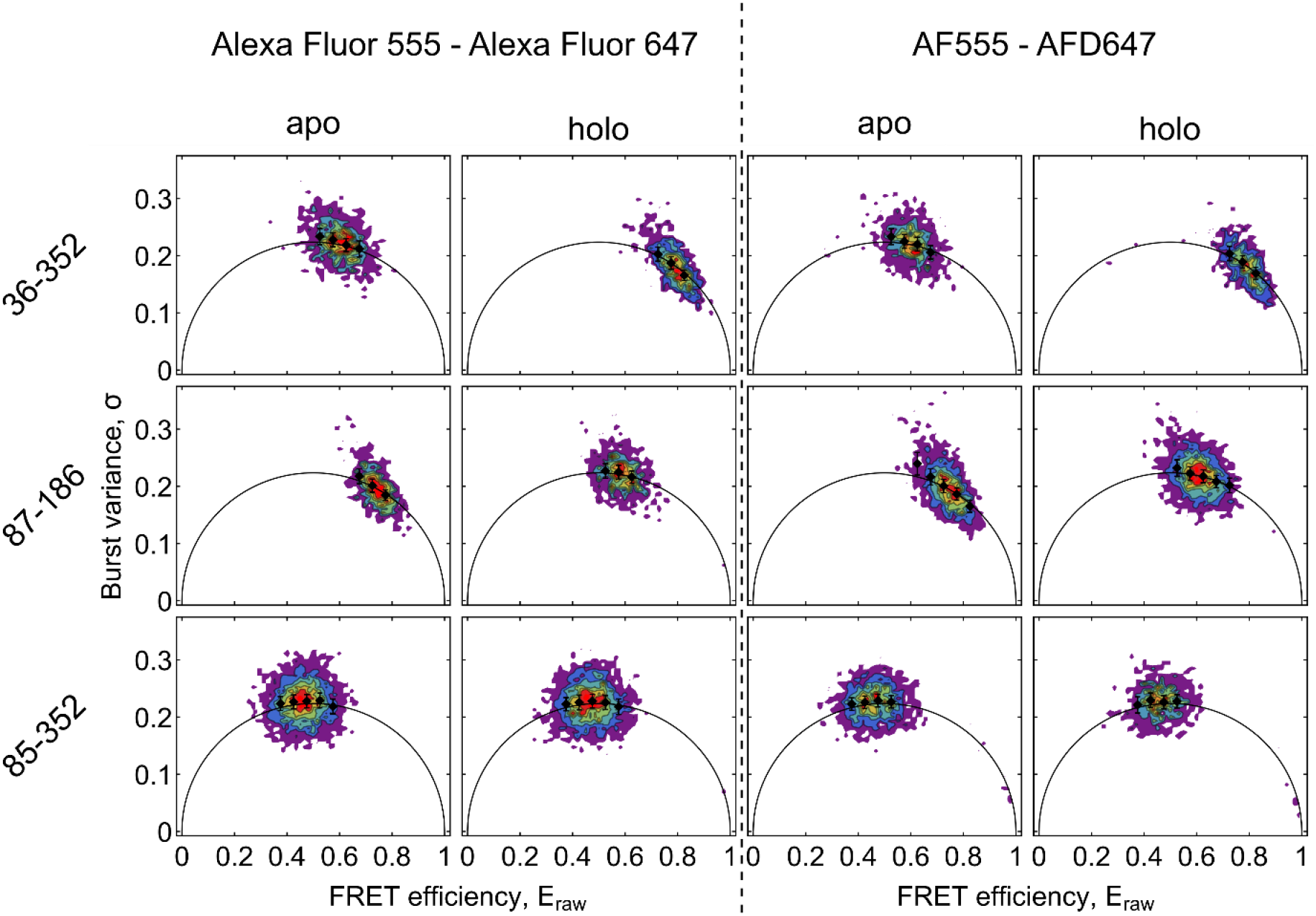
Burst variance analysis. Burst variance analysis of all FRET examples in their apo and holo states for the Alexa-fluorophore-pair (left) and the AF-fluorophore-pair (right). Data are binned into bins of 0.05 and mean and standard error of mean are shown (black).

**Fig. S10.**
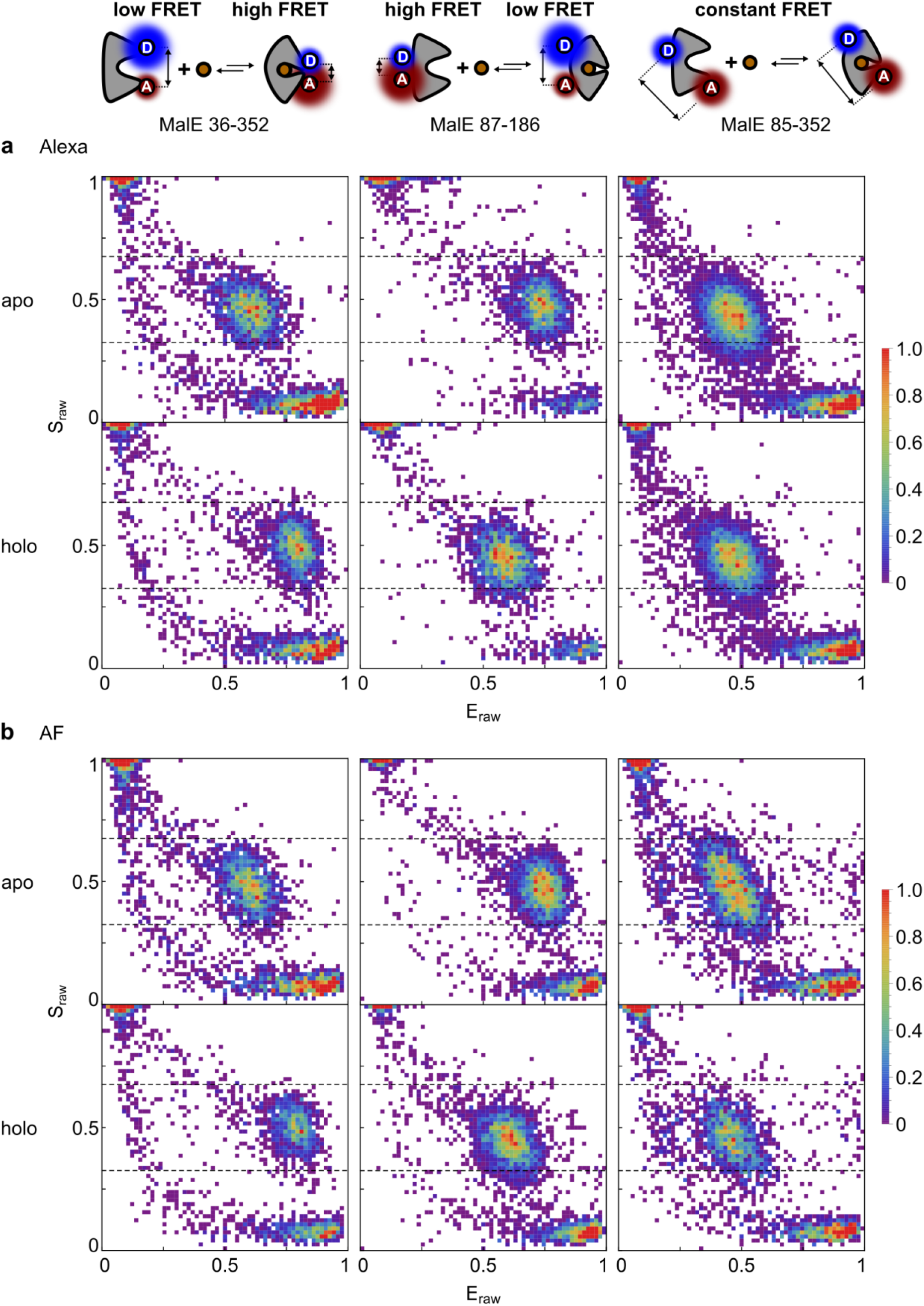
ES-histograms of smFRET experiments. **(a)** ES-histograms from all-photon burst search (threshold 150 photons) of all FRET mutatns in their apo and holo states for the Alexa-fluorophore-pair and **(b)** the AF-fluorophore-pair. Data are binned into 61 bins.

**Fig. S11.**
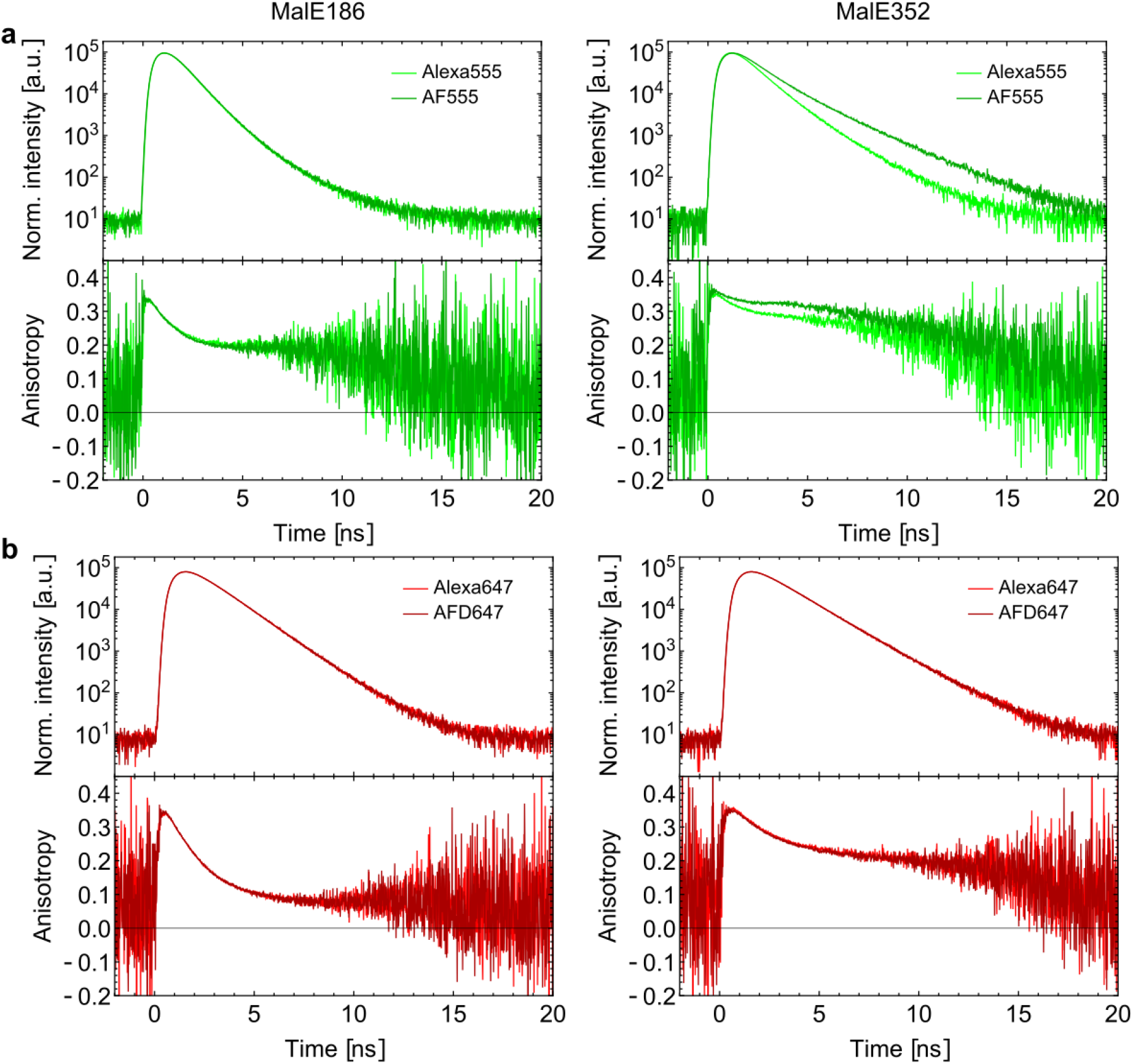
Fluorescent characterization of labelled proteins in bound state: **(a)** Lifetime (top) and time-resolved anisotropy measurements (bottom) of Alexa Fluor Alexa 555 (lighter green) and AF555 (darker green) labelled at residues 186 (left) and 352 (right) in bound state of MalE with 1 mM maltose in buffer. **(b)** Lifetime (top) and time-resolved anisotropy measurements (bottom) of Alexa Fluor Alexa 647 (lighter red) and AF647 (darker red) labelled at residues 186 (left) and 352 (right) in unbound state.

**Fig. S12:**
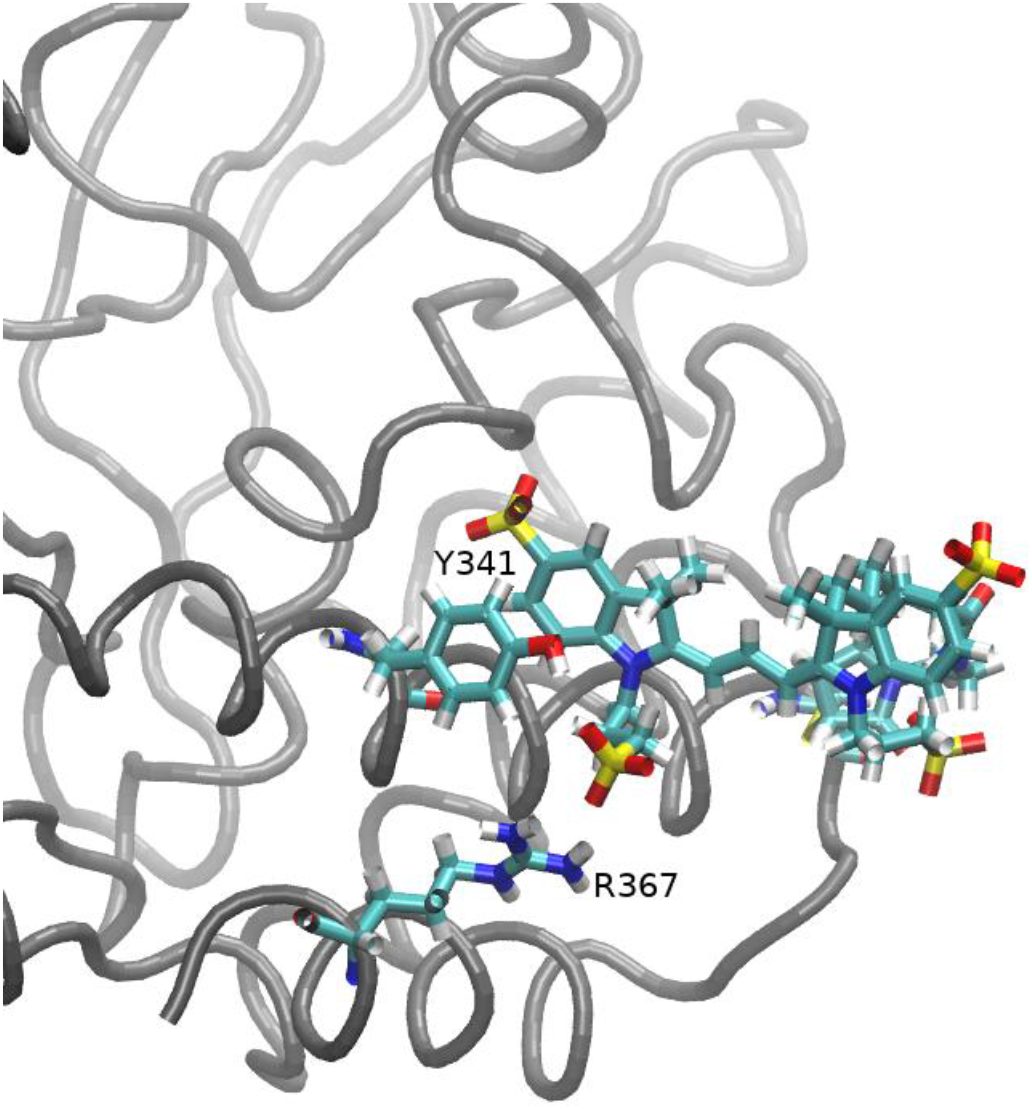
Fluorophore Alexa Fluor 555 #1 attached to mutant S352C of the maltose binding protein. Residues Y341 and R367 as well as the fluorophore are highlighted in stick representation. The depicted picture is a representative snapshot sampled during a 200 ns simulation throughout which frequent hydrogen bond formation (occurrence > 20%) between the oxygen atoms of the SO_3_^-^ group attached to the five-membered ring of the second fluorophore indole ring and the OH group of Y341 or the NH1 and NH2 groups of R367 occurs. The sampled distances between the involved acceptor oxygen atoms and the donor atoms of the amino acids are shown in Fig. S13. In comparison, none of the performed simulations of fluorophores attached to the A186C site presents a hydrogen bond with an occurrence > 20%, involving the oxygen atoms of a SO_3_^-^ group. Moreover, it was found that at the A186C site, Alexa Fluor 555 #1 has significantly less hydrogen bonding interactions with the protein than Alexa Fluor 555 #2 and AF555 (data not shown).

**Fig. S13:**
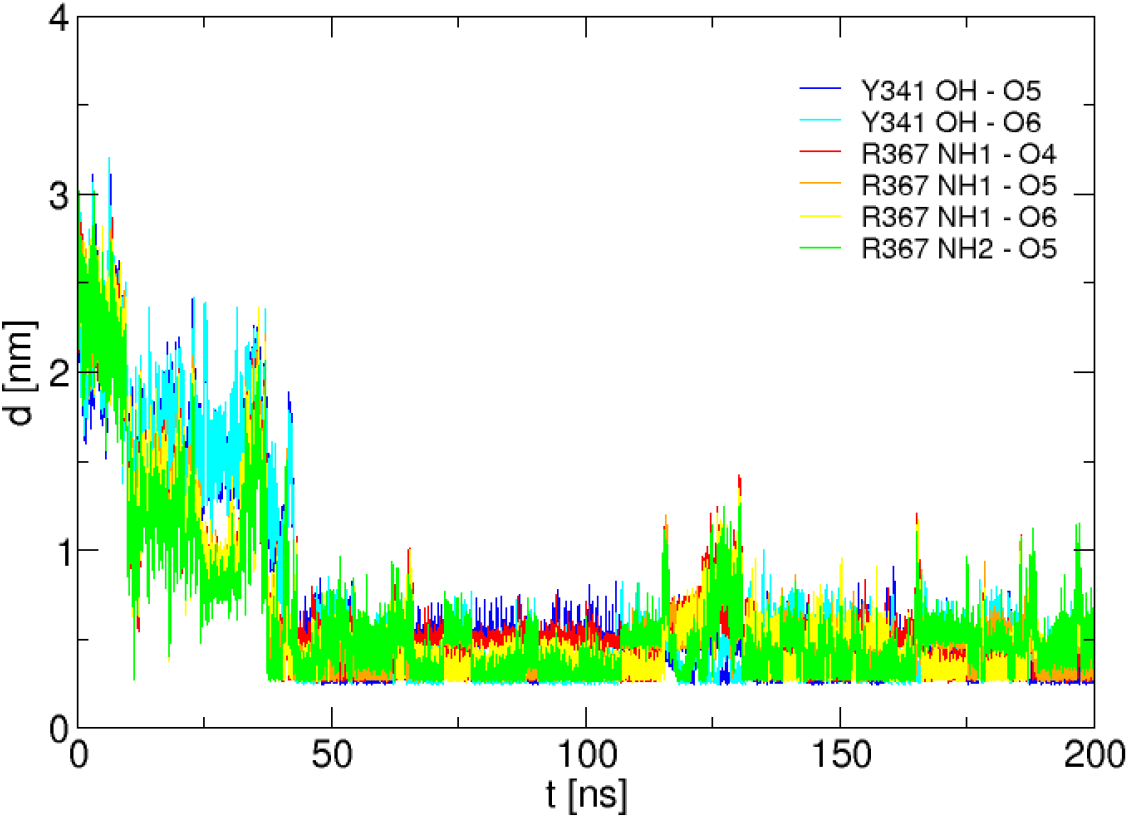
Distances between selected atoms. Distances between oxygen atoms (O4, O5 or O6) of the SO_3_^-^ group attached to the five-membered ring of the second fluorophore indole ring and the oxygen atom of the OH group of Y341 or the nitrogen atoms of the NH1 and NH2 groups of R367 sampled during a 200 ns simulation of fluorophore Alexa Fluor 555 #1 attached to mutant S352C of the maltose binding protein.

**Fig. S14:**
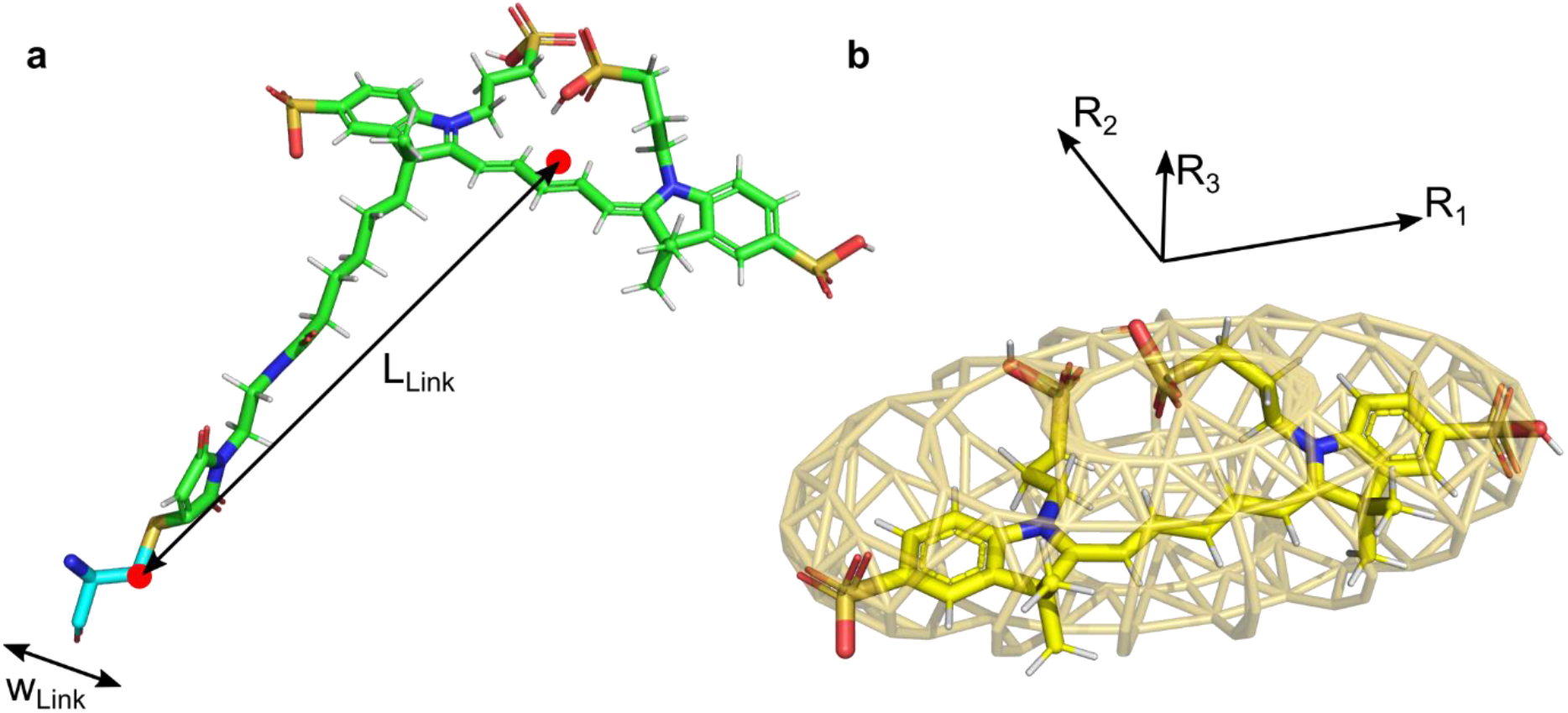
Fluorophore parametrization for FPS with Alexa Fluor 647. **(a)** The linker width is set to be 4.5 Å as suggested for comparable linkers^117^. The linker length is measured between the C-Beta atom and the central of the fluorophore core. (b) The fluorophore core is approximated by an ellipsoid with three radii R1, R2, and R3.

## Notes

### Competing Interest Statement

The authors have declared no competing interest.

